# Mechanisms of tissue-specific genetic regulation revealed by latent factors across eQTLs

**DOI:** 10.1101/785584

**Authors:** Yuan He, Surya B. Chhetri, Marios Arvanitis, Kaushik Srinivasan, François Aguet, Kristin G. Ardlie, Alvaro N. Barbeira, Rodrigo Bonazzola, Hae Kyung Im, GTEx Consortium, Christopher D. Brown, Alexis Battle

**Affiliations:** Department of Biomedical Engineering, Johns Hopkins University, 21218 Baltimore, MD, USA; HudsonAlpha Institute for Biotechnology, 35806 Huntsville, AL, USA; Department of Medicine, Division of Cardiology, Johns Hopkins University, 21287 Baltimore, MD, USA; Department of Computer Science, Johns Hopkins University, 21218 Baltimore, MD, USA; The Broad Institute of MIT and Harvard, Cambridge, MA, USA; Section of Genetic Medicine, Department of Medicine, The University of Chicago, Chicago, IL, USA; Department of Genetics, Perelman School of Medicine, University of Pennsylvania, 19104 Philadelphia, PA, USA

**Keywords:** Matrix factorization, Universal eQTLs, Tissue-specific eQTLs, Transcription factors

## Abstract

**Background:** Genetic regulation of gene expression, revealed by expression quantitative trait loci (eQTLs), varies across tissues in complex patterns ranging from highly tissue-specific effects to effects shared across many or all tissues. Improved characterization of these patterns may allow us to better understand the biological mechanisms that underlie tissue-specific gene regulation and disease etiology.

**Results:** We develop a constrained matrix factorization model to learn patterns of tissue sharing and tissue specificity of eQTLs across 49 human tissues from the Genotype-Tissue Expression (GTEx) project. The learned factors include patterns reflecting tissues with known biological similarity or shared cell types, in addition to a dense factor representing a universal genetic effect across all tissues. To explore the regulatory mechanisms that generate tissue-specific patterns of expression, we evaluate chromatin state enrichment and identify specific transcription factors with binding sites enriched for eQTLs from each factor.

**Conclusions:** Our results demonstrate that matrix factorization can be applied to learn the tissue specificity pattern of eQTLs and that it exhibits better biological interpretability than heuristic methods. We present a framework to characterize the tissue specificity of eQTLs, and we identify examples of tissue-specific eQTLs that may be driven by tissue-specific transcription factor (TF) binding, with relevance to interpretation of disease association.

## Background

Understanding the genetic effects on gene expression is essential to characterizing the gene regulatory landscape and provides insights into the molecular basis of phenotypes. Expression quantitative trait locus (eQTL) studies using genotype and gene expression data have demonstrated that the genetic regulation of gene expression is pervasive [1, 2, 3, 4, 5]. Additionally, numerous studies have leveraged eQTLs to characterize the molecular basis of complex phenotypic variation [6, 7, 8, 9, 10].

Tissues in the human body carry out universal cellular processes in addition to performing highly specialized functions, driven in large part by patterns of gene expression in each cell type. Characterizing the tissue sharing and tissue specificity of genetic effects on gene expression is therefore critical to understanding how genetic variation leads to phenotypic changes. Recent work has identified eQTLs across a broad range of human tissues. The Genotype-Tissue Expression (GTEx) project has collected eQTL data across 49 human tissues (Additional file 1: Figure S1), which provide an unprecedented opportunity to uncover the universal and tissue-specific patterns of genetic regulation of gene expression [1].

Several methods have been developed to capture the underlying tissue-specific architecture in eQTLs across tissues. The simplest such method is based on the effect sizes or P values of eQTLs to identify eQTLs specific to individual tissues or cell types [11, 12]. This method, while easily implemented, requires subjective thresholds and ignores the underlying similarity of tissues. Statistical frameworks have been developed to jointly analyze eQTLs from different datasets, such as eQTL-BMA and Meta-Tissue [13, 14]. These methods are more computationally demanding but potentially more accurate in their estimation of tissue specificity. However, neither class of method addresses the inherent patterns of similarity of multiple tissues in datasets such as GTEx, which may be relevant to patterns of shared mechanism.

For example, a regulatory mechanism relevant to endothelium would affect several related tissues, but would not be universal across all tissues. Evaluating functional properties of eQTLs that are highly specific only to the stomach would exclude those shared across endothelial tissues, and thus would not reveal the regulatory mechanisms shared across endothelial tissues. Manually identifying relevant groupings of tissues is not always obvious or feasible, and, furthermore, such groupings do not form mutually exclusive sets of tissues. Matrix factorization applied to eQTL statistics offers a more flexible and natural approach for identifying underlying patterns across eQTLs that may indeed better reflect biological mechanisms which likewise act across related, non-mutually exclusive subsets of tissues or samples [15]. Recently, matrix factorization has been applied in a Bayesian setting to capture the structure of genetic regulation in human tissues, however specific modelling choices for factorizing eQTL effects in various domains remain to be comprehensively evaluated [16]. It is further unexplored what insights into regulatory mechanism and functional consequences can be gained by evaluating these complex patterns of universal and tissue-specific eQTL effects.

In this study, we propose a constrained matrix factorization model called weighted semi-nonnegative sparse matrix factorization (sn-spMF) and apply it to analyze eQTLs across 49 human tissues from the GTEx consortium. We learn a lower-dimensional representation of eQTL effects across tissues, capturing both tissue-shared and tissue-specific patterns of eQTL activity. We leverage this atlas of universal and tissue-specific eQTLs to begin to characterize the regulatory mechanisms that underlie this specificity, and compare this approach to standard methods of identifying tissue-specific eQTLs. We demonstrate that the universal and tissue-specific eQTLs exhibit distinct patterns of cis-regulatory element enrichment and identify specific TFs that appear to drive tissue-specific genetic effects.

## Results

### Matrix factorization of multi-tissue eQTL effects

The effect of eQTL variants on gene expression varies across tissues, as has been previously observed [1, 2, 17]. To better understand common patterns of genetic impact across tissues and to characterize the mechanisms that underlie tissue specificity, we developed and applied a matrix factorization model called semi-nonnegative sparse matrix factorization (sn-spMF). This model assumes that the effect of an eQTL across tissues is a linear combination of “factors”, where every factor represents a common pattern of eQTL sharing across particular sets of tissues (Fig. 1A). Then, for a given eQTL, the loadings, or “weights,” on each factor reflect how strongly that eQTL’s effects are explained by that factor (and corresponding tissues). Given a multi-tissue dataset of eQTL association statistics as input, we identified a set of explanatory tissue factors by minimizing an objective function combining two components: (1) a weighted squared error term that captures how well the learned weights and factors reconstruct the observed eQTL effect sizes and (2) a regularization term that encourages sparsity in both factors and weights through an L1 penalty (Fig. 1B). Since it has previously been shown that inconsistent directions of effect for eQTLs will often arise from allelic heterogeneity rather than true sharing [18], we constrained factors to be nonnegative.

**Figure 1.**
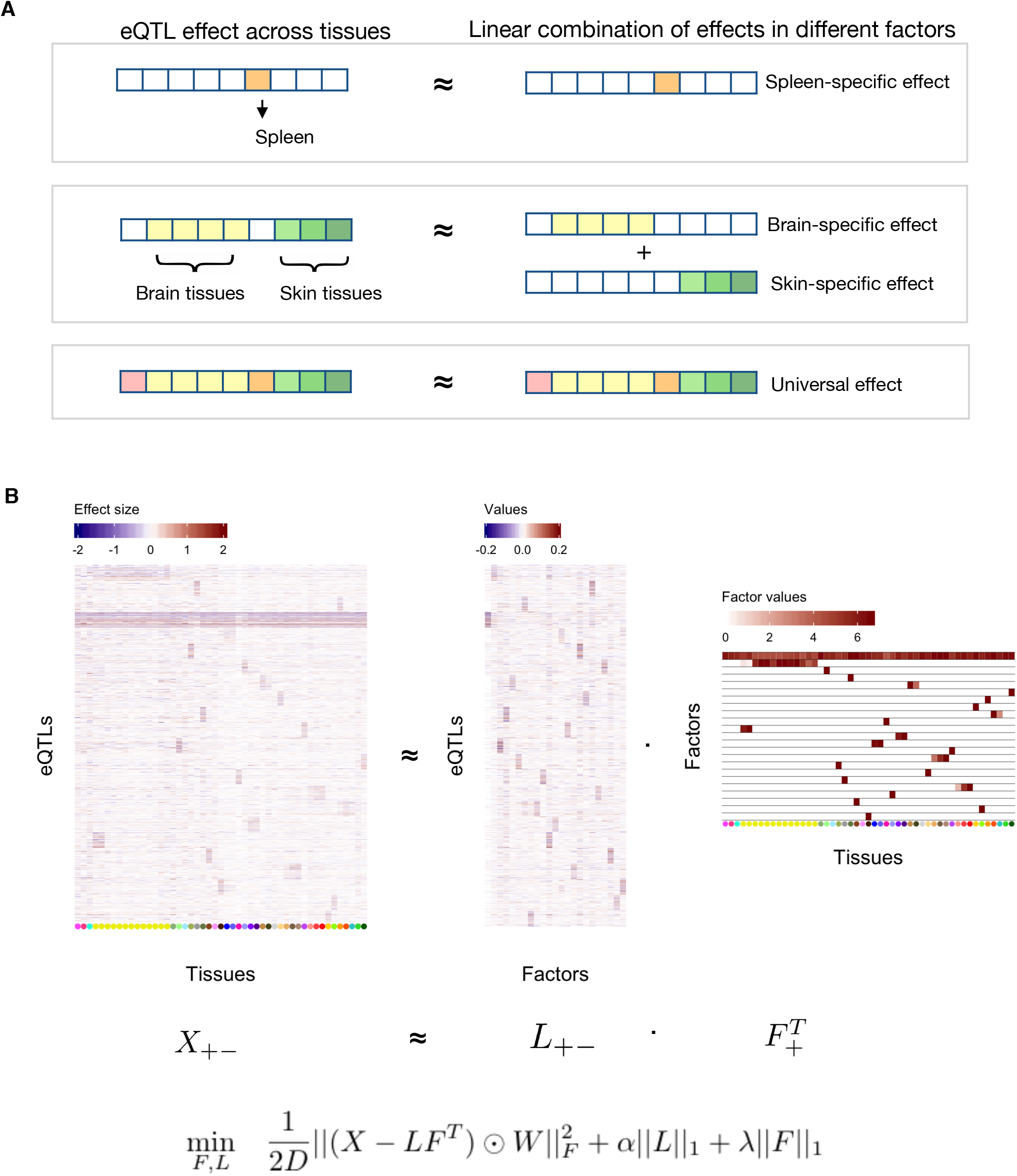
Matrix factorization model to dissect eQTL effects across tissues. **A** Simplified example of the relationship between eQTL effect sizes and factors. Upper panel: The effect of an eQTL in spleen can be represented by a spleen specific factor. Middle panel: The effect of an eQTL in four brain tissues and three skin tissues can be summarized as the summation of brain-specific effect and skin-specific effect. Lower panel: The effect of an eQTL in all nine tissues can be summarized as a universal effect across all tissues. **B** Learning factors underlying eQTL effects from GTEx. X matrix represents the effect size of eQTLs across tissues. For visualization, a subset of the eQTLs are shown. Patterns of tissue sharing and tissue specificity are observed in X. Matrix factorization is implemented to learn the factor matrix *F*, where each factor captures a pattern of eQTL effect sizes across the tissues. GTEx tissues are color-coded for reference (Additional File 1: Figure S1)

By optimizing the objective function using alternating least squares applied to the GTEx v8 data across 49 tissues, we learned a factor matrix *F* with 23 factors (see Methods, Additional file 1: Figure S1, S2). These factors can be categorized into two major types: a universal factor, which captures eQTLs with largely consistent effects across all 49 tissues, and tissue-specific factors, which reflect effects only found among subsets of individual tissues. Tissue-specific factors include two subtypes: 8 factors representing combinations of tissues and 14 factors representing single tissues. Each of the 8 multi-tissue factors involves closely related tissues. For example, factor 2 represents effects of eQTLs in 13 brain regions; factor 15 represents effects in transverse colon and small intestine. For interpretability, each factor is named based on the tissues it represents (Additional file 1: Figure S2). In total 41 out of 49 tissues are represented by nonzero values in at least one tissue-specific factor. The 8 tissues that do not appear in any tissue-specific factor have significantly smaller sample sizes compared to the 41 tissues captured by one or more factors (two-sided t-test P value = 0.024), and thus fewer eQTLs are detected that are unique to those tissues.

### Identification of universal and tissue-specific eQTLs

For each individual eQTL, we identified the relevant patterns of tissue sharing by estimating the contribution from each of our learned factors to the eQTL’s effect sizes, using a second pass of weighted linear regression (See Methods). The observed patterns of tissue sharing and how they are decomposed by matrix factorization are illustrated in the four following examples. First, an eQTL for GLT1D1 is highly specific to liver, and loads only on the corresponding liver factor (Fig. 2A). Second, an eQTL for AATF loads on the brain tissues factor and the tibial nerve factor to explain its combined effect size profile (Fig. 2B). Although this eQTL has small effects (or large variance) in some brain sub-regions, the model is able to identify a brain-wide effect as a likely explanatory factor for this eQTL. Third, an eQTL for U2AF1 with relatively consistent effects across tissues loads only on the universal factor (Fig. 2C). Finally, an eQTL for CD14 has consistent effects across all tissues in addition to a stronger effect specific to the testis (Fig. 2D).

**Figure 2.**
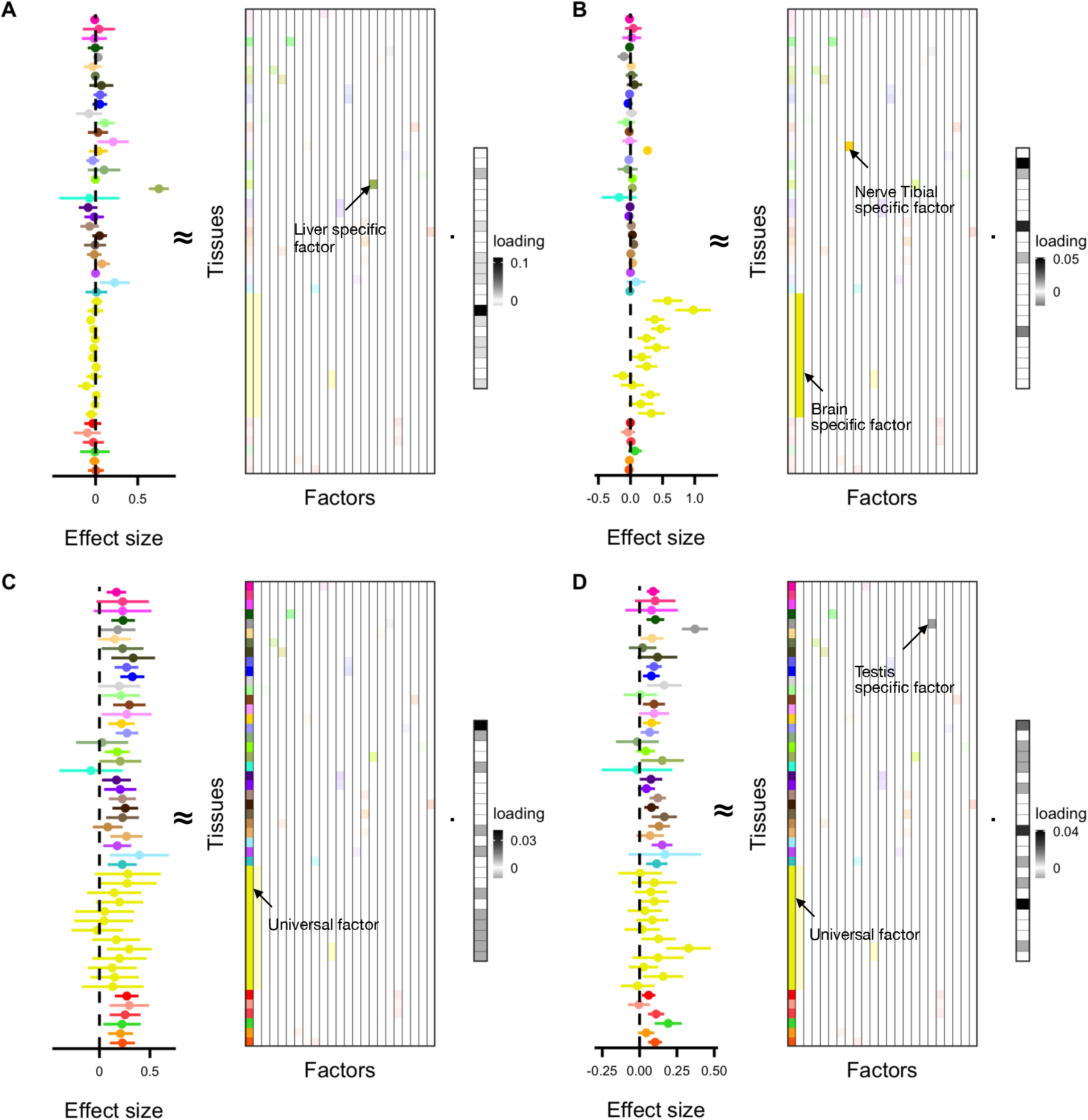
Assignment of eQTLs to factors. In each panel, the effect size and 95% confidence interval of an eQTL across 49 tissues is illustrated. At right, the fitted linear combination of factors is displayed. Factors that have coefficients with FDR >= 0.05 are filled with faded colors. **A** A liver specific eQTL (GLT1D1 - rs1012994). **B** An eQTL (AATF - rs76014915) with activity in brain tissues and tibial nerve. **C** A universal eQTL (U2AF1 - rs234719). **D** An eQTL (CD14 - rs2563249) with universal and testis specific effects.

In summary, 1,076,761 eQTLs (20% of tested eQTLs) load on the universal factor; we refer to these eQTLs as “universal eQTLs” (u-eQTLs). For each tissue-specific factor, 76, 976 to 431, 585 eQTLs (1.5% to 8.1% of tested eQTLs) have significant loadings; we call these eQTLs “tissue-specific eQTLs” (ts-eQTLs) (Fig. 3A). In total across factors, 2,821,650 eQTLs (53% of tested eQTLs) are found to use at least one tissue-specific factor (Fig. 3B). There are 638,784 eQTLs that load on both the universal factor and tissue-specific factors (59% of the u-eQTLs and 22% of the ts-eQTLs, Fig. 3C), indicating that in addition to a broad, shared effect across tissues, these eQTLs have a much stronger effect on expression in a particular subset of tissues. eQTLs tend to load on a small set of tissue-specific factors, with 3, 083, 103 eQTLs (99% among the eQTLs loaded on at least one factor) using less than six tissue-specific factors (Fig. 3D).

**Figure 3.**
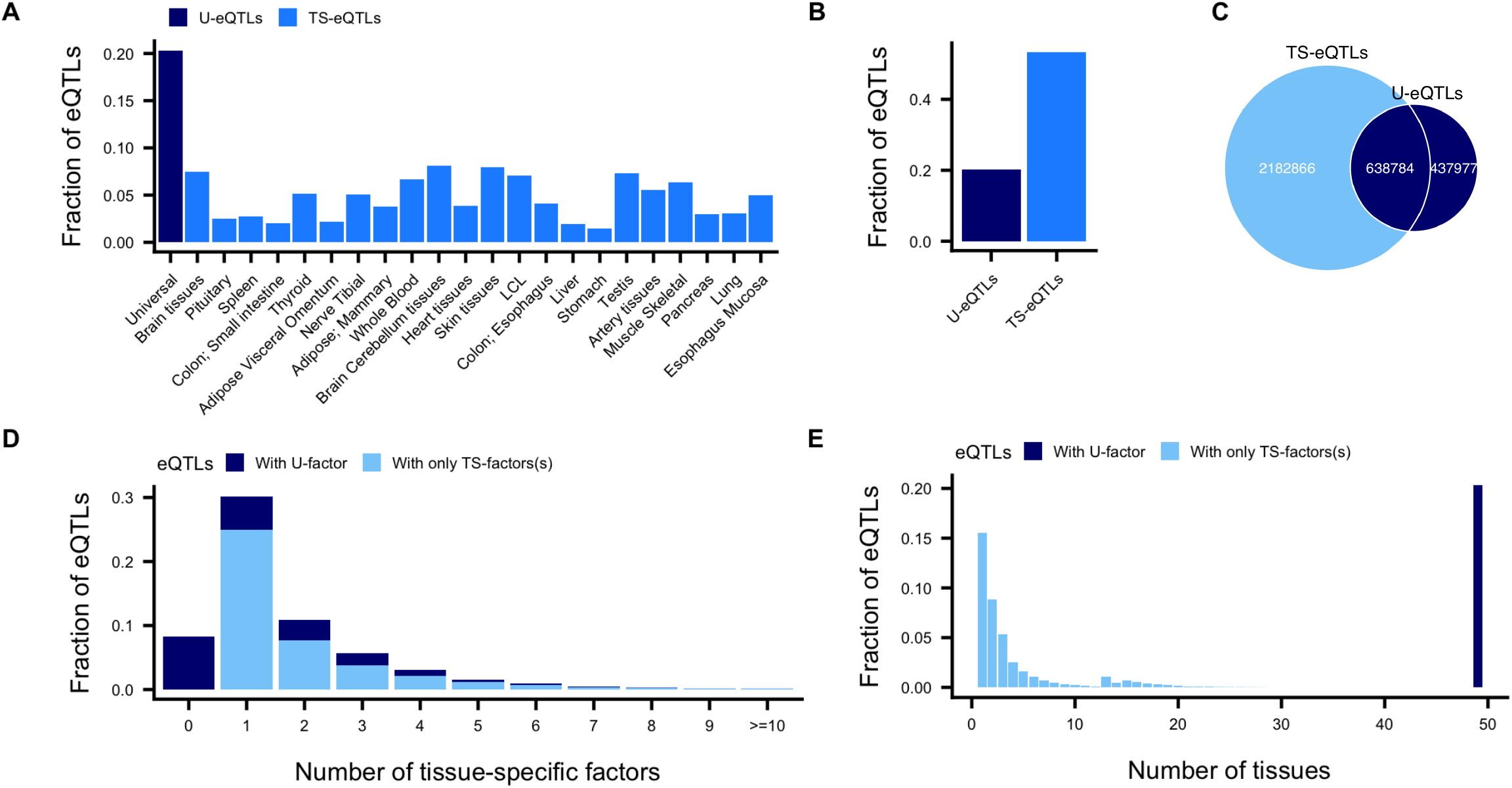
Identification of tissue-specific and universal eQTLs. **A** Fraction of tested eQTLs that load on each factor. **B** Fraction of eQTLs that load on the universal and tissue-specific factors. **C** Number of tested eQTLs that load on the universal factor and overall on any tissue-specific factor (ts-eQTLs). **D** Fraction of eQTLs that load on different number of tissue-specific factors. Because of the uniqueness of the universal factor that is different from the tissue-specific factors, the numbers are shown in two subgroups depend on whether the eQTL has the universal factor. **E** Fraction of eQTLs with activity in different numbers of tissues. The numbers of unique tissues represented in the set of factors for each eQTL are summed.

The number of factors an eQTL loads on should provide a more biologically interpretable indication of the number of independent contexts in which an eQTL is active, rather than simply counting individual significant tissues. Datasets often contain multiple similar or even duplicate tissues, such as the thirteen brain regions in GTEx, or the two skin tissues that only differ by sun exposure. It may be misleading to count a neuron-specific eQTL as active in thirteen tissues, not at all comparable to a very general eQTL active in thirteen highly distinct tissues. Here, we demonstrate that eQTLs tend to be active in just a few factors, tailing off rapidly, but these factors sometimes correspond to numerous tissues (Fig. 3D, E), providing some interpretation for the familiar “U-shape” curve that has been reported previously ([19], The GTEx Consortium 2019, in submission). However, we note that 8 tissues are not significantly represented by any tissue-specific factor and, therefore, can’t be captured in this analysis.

### Matrix factorization improves biological interpretation over heuristic methods of determining tissue relevance

The method most commonly used to determine ts-eQTLs is simply to apply heuristic thresholds to effect sizes, P values, or meta-analysis results for individual tissues [11, 12, 14, 17]. If an eQTL statistic exceeds the chosen threshold for a given tissue, and remains below another threshold for other tissues, it is considered to be tissue specific. None of these approaches consider the common patterns of tissue sharing and may obscure eQTL mechanisms shared across a subset of tissues (such as brain or endothelium) that were not manually predefined for investigation.

Based on heuristic thresholding on individual tissue P values (see Methods), we identified 312, 502 u-eQTLs and between 1, 374 and 102, 414 ts-eQTLs per tissue – far fewer eQTLs are confidently assigned to each category compared to results from sn-spMF (Additional file 1: Figure S3). This difference is partly because thresholding allows only one pattern (a single tissue or a universal effect) to be assigned to each eQTL, while matrix factorization allows multiple factors and tissues to be involved in explaining the effect size of an eQTL. In addition, thresholding often misses small effects from similar tissues, while matrix factorization is able to aggregate effects for similar tissues. In subsequent sections, we show that matrix factorization allows for the identification of more biologically coherent eQTLs than heuristic approaches do.

#### Tissue-specific eQTL gene function

To examine the functional relevance of ts-eQTL genes, we ran enrichment analysis using biological processes from the Gene Ontology (GO) project [20]. We first evaluated genes with ts-eQTLs and no u-eQTL. For sn-spMF, these eQTL genes are enriched for 546 unique GO terms at FDR < 0.05 (Additional file 1: Figure S4), and the top enriched GO terms are relevant to the corresponding tissues (Additional file 1: Figure S5, S6, S7). The ts-eQTL genes from the standard thresholding method, however, are less enriched in GO biological processes (110 enriched at FDR < 0.05, Additional file 1: Figure S8).

After initial enrichment analysis, we used a more stringent definition of tissue-specificity to restrict the analysis to the genes most unique to each factor. For sn-spMF, we selected genes appearing in less than 6 tissue-specific factors (on average 252 genes per factor). 63 unique GO terms are enriched at FDR < 0.1. The enriched GO terms are related to the matched tissue(s) of the eQTLs (Fig. 4). For example, five GO terms are enriched among liver-specific genes including four metabolic processes (for steroid, drug, uronic acid, and flavonoid) and response to xenobiotic stimulus, each relevant to liver function. For the heuristic method, we selected genes appearing in less than 7 tissues (on average 325 genes per tissue) such that the gene sets are of comparable sizes. No GO term is enriched among these gene sets (Additional file 1: Figure S8). These results indicate that sn-spMF is able to identify eQTL genes with biological functions relevant in the corresponding tissues more effectively than heuristic methods, even with comparably stringent definitions of tissue-specific eQTL genes providing similar numbers of genes for analysis.

**Figure 4.**
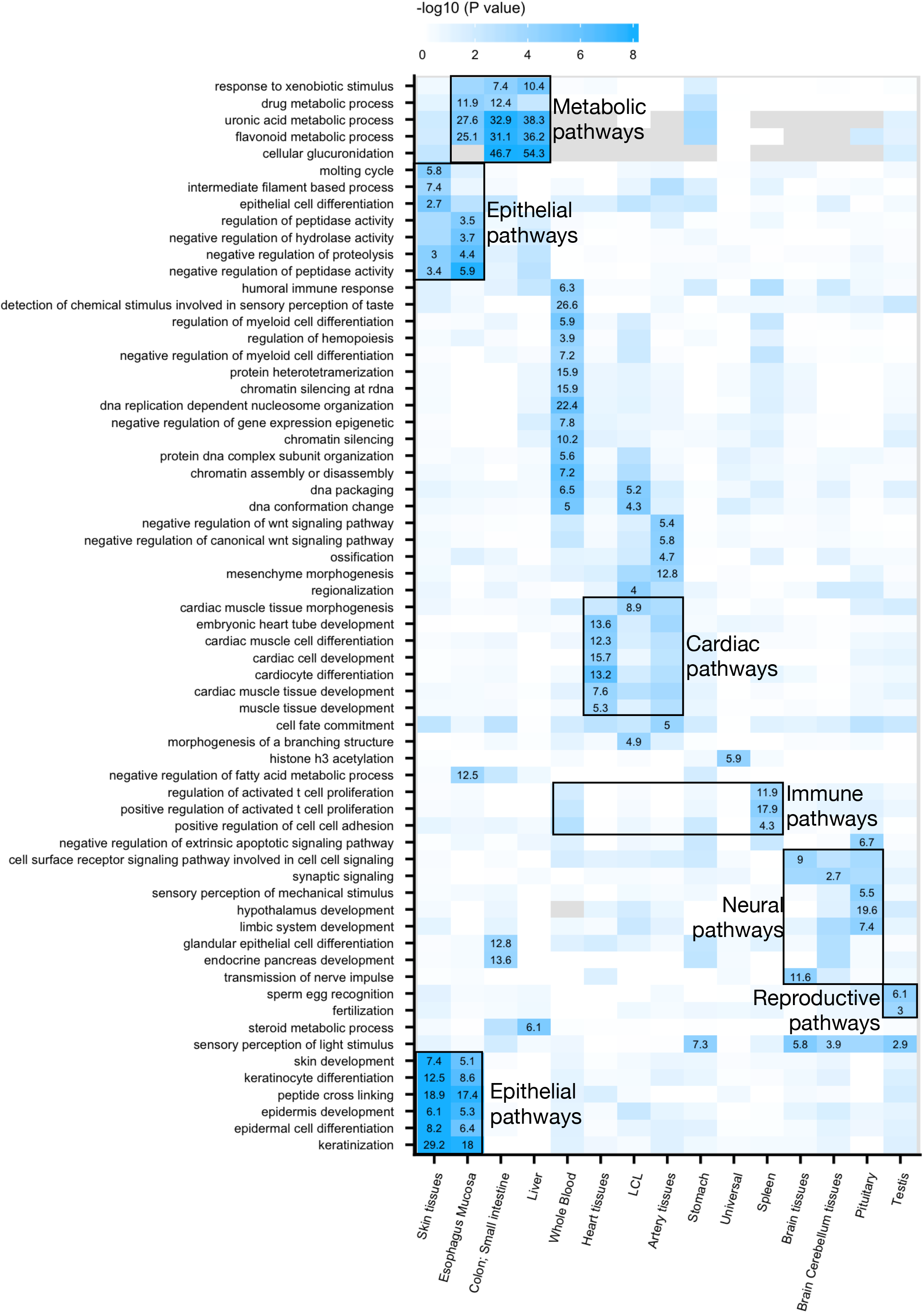
Enriched GO terms for eQTL genes from sn-spMF at FDR < 0.1. Color represents the level of enrichment (– log 10 P value). The GO terms that are significantly enriched are annotated by numbers that represent the odds ratio. GO terms and factors are ordered by hierarchical clustering. Examples of relevant GO terms in related tissues are annotated.

#### eQTL variant enrichment in cis-regulatory regions

eQTL variants are enriched in cis-regulatory elements, including cell-type-specific promoters and enhancers [1, 21, 22]. Consistent with prior observations, u-eQTL variants identified by sn-spMF are more enriched in promoters (OR = 1.9, P value < 2.2 × 10^-16^) than ts-eQTL variants (OR = 1.6, P value < 2.2 × 10^-16^), while ts-eQTL variants are more strongly enriched in enhancers (OR = 1.3, P value = 1.6 × 10^-10^) than u-eQTL variants (OR = 1.0, P value = 0.40, Additional file 1: Figure S9) [1, 23, 24]. Compared to sn-spMF, heuristically defined ts-eQTLs exhibit comparable enrichment magnitude in enhancers (OR = 1.3, P value = 1.2× 10^-6^), but sn-spMF provides an order of magnitude more ts-eQTLs (Additional file 1: Figure S3, S9). While heuristic methods identify highly tissue-specific eQTLs by selecting those with effects clearly limited to a single tissue, sn-spMF identifies many more eQTLs relevant to each tissue-specific factor, each related to a shared set of cis-regulatory elements.

#### eQTL enrichment in transcription factor binding sites

To systematically assess whether eQTLs for each factor are enriched in binding sites for specific TFs, we performed enrichment analysis for each of the 579 TF motifs available in the JASPAR database [25]. As a proxy for TF binding sites (TFBS) in individual tissues, we identified TF motif instances overlapping predicted enhancers and promoters [26, 27, 28, 29].

Enrichment analysis was performed separately for TFBS in promoters and TFBS in enhancers (Methods). In promoters, u-eQTLs and ts-eQTLs are enriched for TFBS of 147 and 185 unique TFs (median = 22 across factors), respectively (FDR < 0.05, Fig. 5A, B). In enhancers, u-eQTLs and ts-eQTLs are enriched for TFBS of 22 and 265 unique TFs (median = 43 across factors), respectively (FDR < 0.05, Fig. 5A, B). Among these 265 TFs, 246 (93%) are enriched for fewer than six tissue-specific factors (Fig. 5C). 0% – 23% (among factors, median 8%) TFs are enriched in both promoters and enhancers (Additional file 1: Figure S10). These results indicate that ts-eQTLs are more enriched in binding sites of particular TFs in enhancers than promoters, while u-eQTLs yield more enrichment in promoters than enhancers. The heuristic approach for identifying ts-eQTLs yields no enrichment of TFBS in promoters and only 11 TFs enriched in enhancers. Similarly, there are fewer TFs enriched for heuristic u-eQTLs (75 in promoters, and 4 in enhancers, Fig. 5A, Additional File 1: Figure S11). The relatively low enrichment of TFBS from heuristically identified eQTLs is presumably due to the much more limited number of eQTLs identified in each category.

**Figure 5.**
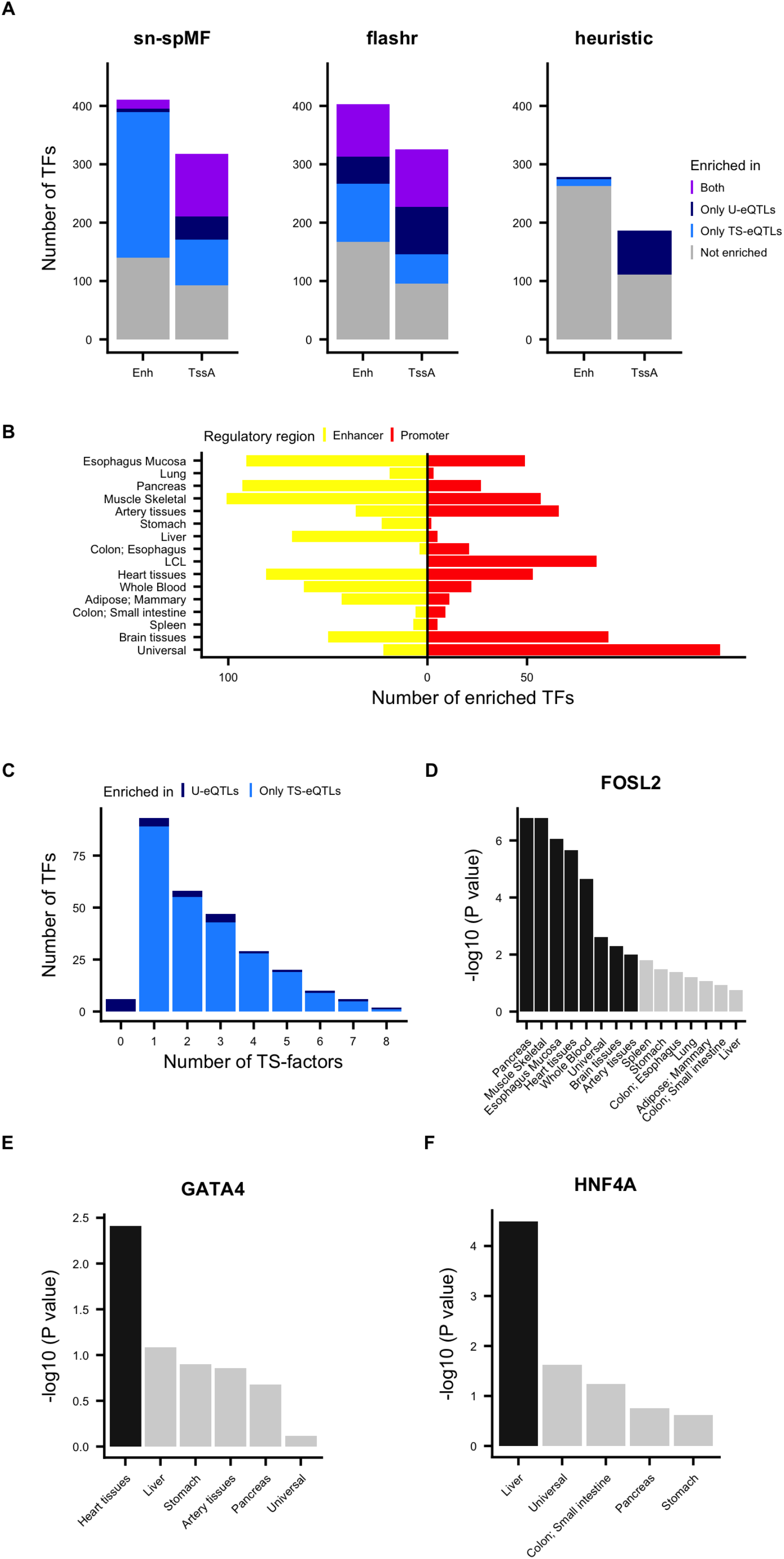
Enrichment of TFBS for u-eQTLs and ts-eQTLs. **A** Number of TFs whose binding sites are enriched for eQTLs across factors at FDR < 0.05 for sn-spMF, flashr, and heuristic methods. Enh - Enhancers, TssA - Active transcription start sites. **B** Total number of TFs with binding sites enriched for either only u-eQTLs, or only ts-eQTLs, or both. **C** Distribution of the number of tissue-specific factors each TF is enriched in. **D, E, F** Enrichment for example TFs among eQTLs across each factor (-log10(P value)) where the TF was expressed in corresponding tissues for (**D**) FOSL2, (**E**) GATA4, (**F**) and HNF4A. Black bars represent that the BH corrected P value is < 0.05.

### Impact of matrix factorization methodological choices

In addition to our sn-spMF model, there are a variety of matrix factorization approaches available. Methodological choices include the selection of priors on loading and factor entries, which may encourage sparsity or other properties, nonnegativity constraints, and hyperparameter selection. One method that has also been applied to eQTL data, flashr, uses a Bayesian framework to automatically learn the sparse structure of effects across tissues, but does not by default impose a nonnegativity constraint on factor values and allows different sparsity penalties to be applied to each factor [16].

To explore matrix factorization choices, we applied flashr to the GTEx data, capturing both universal and sparse factors (Additional file 1: Figure S12) as previously reported [16]. With no nonnegativity constraint, some factors do have mixed signs across tissues, without a clear biological interpretation of this property. Each factor is also somewhat more dense than sn-spMF factors. We used the same second pass linear regression pipeline as in sn-spMF to identify flashr factors relevant to each eQTL. We thus identified 1, 785,127 u-eQTLs and 21, 676 to 487,071 ts-eQTLs (Additional file 1: Figure S13), a comparable number to sn-spMF.

Flashr ts-eQTL genes are comparably enriched for GO biological processes as sn-spMF factors, far exceeding heuristic ts-eQTL genes, with 643 enriched pathways (FDR < 0.05, Additional file 1: Figure S8). However, flashr eQTL variants are not strongly enriched in enhancers (OR = 1.1, Additional file 1: Figure S9). This appears to be due to the denser flashr factors not isolating tissue-specific effects from universal effects as strongly. Assessing TF enrichment, however, because analysis is restricted to enhancers in the relevant tissues, is still able to identify enrichment for 236 TFBS across flashr factors (Fig. 5A, Additional File 1: Figure S14). While regulatory element enrichment appears sensitive to matrix factorization methodological choices, both versions of matrix factorization show advantages over heuristic approaches for identifying tissue-relevant eQTL genes, and for identifying particular transcription factors whose binding sites are impacted by ts-eQTL variants.

### Transcription factors enriched in u-eQTLs and ts-eQTLs

Given the limited systematic research on the consequences of genetic variation within tissue-specific TFBS, we examined the characteristics of TFBS enriched in ts-eQTLs for each factor and in u-eQTLs. We focused on the TFBS found within enhancers because of their generally increased tissue-specific functions (Additional file 1: Figure S9). Binding sites for TFs with broad activity are enriched for u-eQTLs, such as CCAAT/enhancer-binding proteins (CEBPB, CEBPD, CEBPG), T-box 1 (TBX1), AP-1 Transcription Factor Subunit FOSL2 [30, 31, 32, 33] (Fig. 5D). The enrichment of these TFBS in u-eQTLs reflects their participation in a wide range of regulatory processes across tissues.

The enrichment of binding sites for 265 TFs in ts-eQTLs demonstrates their role in regulating gene expression in particular subsets of tissues corresponding to each factor. Among these, binding sites for 176 TFs display enrichment in ts-eQTLs for multiple factors that represent related tissues. For example, hepatic nuclear factor HNF1A, known to be crucial for the development and function of the liver, pancreas and gut epithelium, are enriched for the liver-specific eQTLs, pancreas-specific eQTLs, and ts-eQTLs in colon and small intestine [34, 35]. Furthermore, 89 TFBS are enriched in ts-eQTLs for one tissue-specific factor. Examples include binding sites for the well-characterized cardiac TF GATA4, which are enriched for heart-specific eQTLs [36, 37] (Fig. 5E); hepatocyte nuclear factor HNF4A, which are enriched for liver-specific eQTLs [38, 39] (Fig. 5F); and myogenic factor 4 MYOG, which are enriched for skeletal muscle specific eQTLs [40] (Additional file 1: Figure S15). We continue to explore two TFs in more detail in the following sections. More examples of enriched TFs with previously characterized tissue-specific functions can be found in Additional file 1: Figure S15 and Additional file 2: Table S1.

#### Heart-specific eQTLs are enriched in GATA4 binding sites

Previous studies have demonstrated the essential roles of GATA4 in heart morphogenesis [41]. In mouse studies, GATA4 has been shown to recruit the histone acetyltransferase p300 in a tissue-specific manner in the heart [36]. This GATA4-p300 complex deposits H3K27ac at cardiac enhancers, thus stimulating transcription of genes necessary for heart development. In human, missense mutations in GATA4 are associated with multiple heart diseases such as cardiac septal defects and cardiomyopathy [42, 43]. However, common genetic variants affecting GATA4 TFBS have not previously been shown to be enriched for effects on expression in cardiac tissues. Binding sites of GATA4 in heart enhancers are enriched for heart-specific eQTLs (OR = 1.7, P value = 0.004, Fig. 5E), highlighting the importance of GATA4 in normal physiological conditions of the heart. Among the 48 genes loading on the heart-specific eQTL factor with variants located in TFBS of GATA4, we note that STAT3 has been reported to exhibit a crucial role in cardiomyocyte resistance to physiological stress stimuli [44].

#### Liver-specific eQTLs are enriched in HNFĄA binding sites

Variants in liver-specific HNF4A binding sites are enriched for eQTLs loading on the liver-specific factor (OR = 2.9, P value = 3.3 × 10^-5^, Fig. 6F), which has not been previously shown. HNF4A is an essential TF during liver organogenesis and development [38, 39] and harbors a missense mutation (rs1800961) strongly associated with liver relevant traits including high-density lipoprotein levels and total cholesterol [45, 46, 47] (Additional file 1: Figure S16).

**Figure 6.**
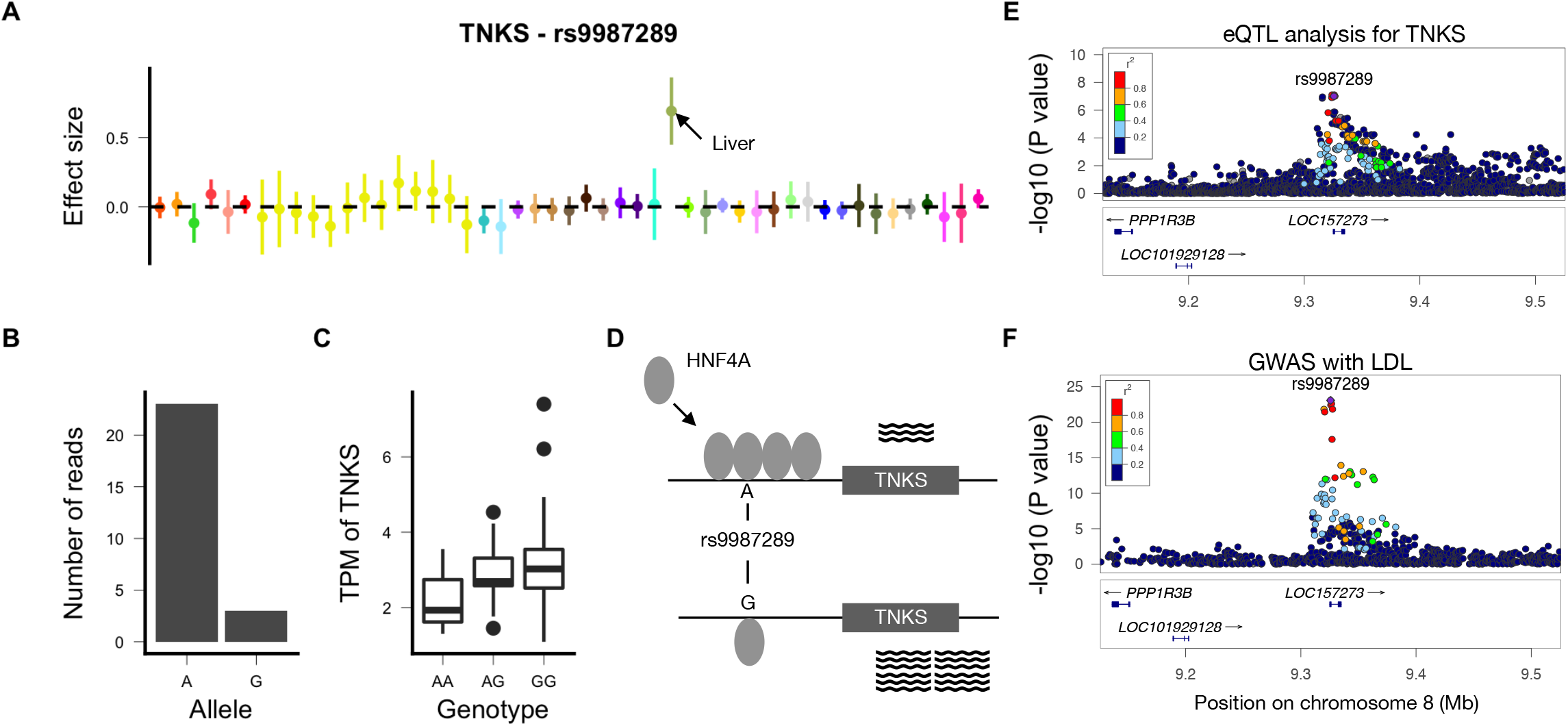
Example liver-specific eQTL, TNKS - rs9987289, in a TFBS of HNF4A and co-localizing with liver-specific phenotypes. **A** Effect size and 95% confidence interval of TNKS - rs9987289 across 49 tissues in GTEx. **B** Allele-specific HNF4A ChIP-seq reads over rs9987289 in the liver (See Methods). **C** Transcripts Per Kilobase Million (TPM) of TNKS in the liver among individuals with different genotypes at rs9987289. **D** Schematic illustration of hypothesized mechanims: allele specific binding of HNF4A at rs9987289 and altered levels of expression of TNKS. **E** Manhattan plot (LocusZoom v0.4.8) [69] of TNKS expression levels in the liver around rs9987289. **F** Manhattan plot for LDL GWAS around rs9987289.

With the availability of Chromatin Immunoprecipitation followed by high-throughput Sequencing (ChIP-seq) data for HNF4A in human liver tissues in ENCODE, we are able to directly map the genome-wide binding sites of HNF4A. Replicating the motif-based enrichment described above, liver-specific eQTLs are strongly enriched in HNF4A ChIP-seq peaks (OR = 3.6, P value < 2.2 × 10^-16^). The enrichment is not as strong in ts-eQTLs for other tissues (OR = 1.8 in testis to 2.6 in pancreas). Also, liver-specific eQTLs are significantly more enriched in HNF4A binding sites than are u-eQTLs (OR = 1.7, P value < 2.2 × 10^-16^).

We hypothesized that variants in HNF4A binding sites lead to liver-specific eQTLs via differential binding of HNF4A. We quantified allele-specific binding (ASB) of HNF4A and, as a tissue-shared control, CTCF (see Methods). Liver-specific eQTLs are indeed significantly enriched for ASB of HNF4A (OR = 1.4, P value = 0.003), but not CTCF (OR = 0.8, P value = 0.4). This finding supports the possibility that the enrichment of liver-specific eQTLs in HNF4a motifs reflects altered binding affinity of HNF4A at these eQTL variants, providing a testable hypothesis for experimental validation.

### Example eQTL variant in HNF4A binding site relevant to liver phenotypes

Among the liver-specific eQTLs identified by sn-spMF, rs9987289 exhibits significant ASB for HNF4A (Fig. 6A,B, Additional file 1: Figure S17). The A allele is associated with increased HNF4A binding (ChIP-seq read ratio = 7.7, two-tailed binomial test P value = 8.8 × 10^-5^) and with significantly lower expression of the eGene TNKS (Fig. 6B, C). HNF4A may act as a repressor of TNKS, and these data suggest that the A allele of rs9987289 may act by increasing binding of HNF4A and therefore reducing expression levels of TNKS. Though HNF4A has been widely reported as a transcriptional activator, it has also been associated with transcriptional repression [48, 49, 50, 51, 52] (Fig. 6D). Rs9987289 is located in a flanking active promoter (TssAFlank) region surrounded by enhancers in liver, while it is found in quiescent or heterochromatin regions in all 13 non-liver tissues where HNF4A is expressed (Additional file 1: Figure S18, S19).

Furthermore, rs9987289 is significantly associated with several liver-related phenotypes, including low-density lipoproteins (LDL) cholesterol levels and high-density lipoproteins (HDL) cholesterol levels [REF GTEx GWAS companion] [45] (Additional file 1: Figure S20). The liver eQTL of TNKS and the association statistics for LDL are strongly co-localized (posterior probability of rs9987289 being causal for the shared signal = 0.64) [53] (Fig. 6E, F). Though TNKS has been widely recognized for its role in controlling telomere length, there is emerging evidence of TNKS participating in liver metabolism [54, 55].

Together, these results support the hypothesis that the tissue-specific regulatory effect of ts-eQTL variant rs9987289 in liver may have phenotypic consequences: an active cis-regulatory element unique to liver, allele-specific binding of liver TF HNF4A in hepatocytes, and finally co-localization of the eQTL effect with lipid GWAS hit. Such examples can provide testable hypotheses regarding multiple steps of the mechanism through which genetic variation may affect a high-level phenotype.

## Discussion and Conclusions

In this study, we explored the genomic context and potential mechanisms underlying tissue-specific effects of genetic variation by applying a constrained matrix factorization model (sn-spMF) to multi-tissue eQTL data from the GTEx project. Using sn-spMF, we learned factors representing the common patterns of eQTL sharing across tissues, such as factors corresponding to universal effects across all tissues and effects shared among only brain tissues or among muscle tissues. This allowed us to explore eQTL effects shared across overlapping subsets of tissues that share cis-regulatory mechanisms due to shared cell types or developmental origin, without having to manually prespecify each such pattern. These learned factors enabled us to evaluate potential mechanisms relevant to genetic effects following these patterns of tissue sharing.

sn-spMF identified much larger sets of tissue-specific eQTLs than did heuristic methods. The ts-eQTLs from sn-spMF were also equally or more enriched for GO biological processes, transcription factor binding sites, and tissue-specific cis-regulatory elements than the heuristic ts-eQTLs. These results suggest that sn-spMF identifies larger numbers of ts-eQTLs that remain biologically coherent, offering an opportunity for novel mechanistic insights. Other versions of matrix factorization, such as flashr, also provide meaningful views of tissue specificity.

The large set of ts-eQTLs provided by sn-spMF enabled a detailed evaluation of eQTLs in transcription factor binding sites that was not possible from heuristic approaches. We evaluated 76, 976 to 431, 585 ts-eQTLs for enrichment in promoter and enhancer elements, and were able to identify 185 and 265 TFs enriched among these, respectively. This list of 265 TFs enriched in ts-eQTL enhancers provides experimentally testable hypotheses about specific genetic variants within TFBS that alter expression in a tissue-specific fashion.

Matrix factorization is inherently limited by the eQTL data used as input to the method – any tissue that is underpowered or not well represented in the original eQTL dataset is unlikely to be captured strongly by a ts-eQTL factor with sn-spMF. Further, sn-spMF does not explicitly model linkage disequilibrium (LD) or consider allelic heterogeneity, rather it relies on the user to pre-select candidate causal variants using fine-mapping tools or other approaches. Additionally, many matrix factorization approaches, priors, and constraints remain to be explored that may capture different properties of the eQTL data than represented here. Different applications, such as time series or perturbation-response eQTL data may ultimately benefit from specialized matrix factorization formulations [15].

In conclusion, we have developed a constrained matrix factorization model to learn patterns of eQTL tissue specificity across 49 human tissues using data from GTEx v8. We observed improved enrichment of biologically relevant genes and cis-regulatory elements compared to heuristic methods. Matrix factorization also revealed the potential impact of ubiquitous TFs on universal eQTLs and provided a list of candidate TFs relevant to each tissue-specific set of eQTLs.

## Methods

### Data

GTEx Release v8 project has collected both genotype data from whole genome sequencing (WGS) and RNA sequence (RNA-seq) data of 15, 253 samples, consisting of 47 tissues and two cell lines from 838 individuals (The GTEx Consortium 2019, in submission). GTEx v8 data release includes cis-eQTL analyses that test for association between gene expression and variants within 1MB of the genes’ transcription start sites (TSS). To restrict the analysis to potential casual variants, we used cis-eQTLs that are in the 95% credible set for at least one tissue [56]. That is to say, for each eQTL gene, the credible set consists of eQTL variants that include the causal variant with 95% probability. In total, 5, 301, 827 eQTLs with 17, 480 unique protein coding eQTL genes are included in the analysis. For these 5, 301, 827 eQTLs, we collected the effect size and standard deviation from univariate cis-eQTL analysis across tissues. Missing data was filled by 0s since it was mostly due to low gene expression in the corresponding tissue and thus little genetic regulation should exist. The lead variants with the most extreme geometric mean P values for the 17, 480 eQTL genes were used as input (matrix *X* and *W*) to learn the factor matrix (matrix *F*).

### Learn the lower dimensional representation of tissues (factor matrix *F*)

eQTL effects across tissues can be represented by a matrix *X_N×T_* where *N* is the number of eQTLs and *T* is the number of tissues. Each row is the effect of an eQTL across all tissues, and each column is the effect of all eQTLs for one tissue. The effect values are in the set R of all real numbers (ie. have mixed signs). The goal is to learn a factor matrix *F_T×K_* such that *X* ≈ *LF*^T^.

### Weighted sparse semi-nonnegative matrix factorization algorithm

In order to describe the eQTL effects, we designed the objective function with several features: 1) Weighted sum of residuals: in order to account for uncertainty in effect size estimates, the residual for each data point was weighted by the reciprocal of its standard error. In this way, the data point with larger effect size contributes more influence over estimating the parameters. 2) Sparseness: to alleviate over-fitting, an l1 penalty was applied to the decomposed matrices. 3) Semi-nonnegativity of the decomposed matrices: the factors capture the pattern of tissues, and thus it was a natural constraint to make the factors nonnegative for ease of interpretation. At the same time, because the input matrix has mixed signs, there was no such constraint on the loading matrix. The factorization can be summarized as: 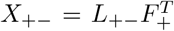. The objective function was formulated as below:

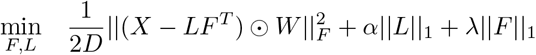

Where *F* is nonnegative, *W* is the element-wise reciprocal of the standard error of eQTL across *M* tissues.

This objective function is biconvex, that is, convex only in *F* or in only *L* given the other. We used alternating least squares (ALS) with gradient descent to solve the problem (Algorithm 1, implemented in `R version 3.5.1`, [57, 58]). At each iteration, we fixed *F* and updated *L*, and then fixed *L* and updated *F*. The update was finished when the Frobenius norm of difference in *F* between two iterations was < 0.01. In each update step the optimization problem was a linear regression with constraints. Since the solution to linear regression was guaranteed to minimize the sum of mean squared error and penalty, the cost function monotonically decreased.

#### Model selection

In the sn-spMF model, we need to decide the rank of the decomposition (*K*) and the sparsity level (*α, λ*). Because of the stochastic nature of matrix factorization, Brunet et al. proposed a method looking for the most stable factorization result, and this method has been applied in various studies [59, 60]. We obtained the consensus matrix *C_M ×m_* after 30 runs with random initialization. The values in *C* are between 0 to 1, representing the proportion of runs in which a pair of tissues are assigned to the same factor. Using the *C* matrix, we computed the cophenetic correlation which is used to measure the degree of dispersion for the *C* matrix. Higher cophenetic correlation indicates more a stable factor matrix. We selected *K* using this criteria. Because factors should be independent from each other to alleviate multicollinearity, we selected the penalty parameters resulting in smallest correlation between the factors.

### Assignment of eQTLs to factors

After we have learned the factors, we mapped the effect of each eQTL to the factors by weighted linear regression. For each eQTL, the weighted linear regression is fit: *x* = *Fl* weighted by its reciprocal of standard error. To alleviate multiple testing burden, we removed the eQTLs those variants in perfect LD (*R*^2^ = 1) with variants from another eQTL before running regression for the remaining 3, 601, 800 eQTLs [61]. Statistical importance is captured by the P values of the factors. We applied Benjamini Hochburg correction to get the corrected P value for every factor [62]. We then mapped the P values back to all 5, 301, 827 eQTLs where the SNPs are in an LD block with the tested SNPs and the genes are the same. We observed that occasionally there were factors assigned negative regression coefficients when the actual observed effect sizes in the corresponding tissues were positive, or vice versa. This discrepancy arose due to colinearity between the factors, and, in such cases, the discrepant factors were not included for downstream analysis. We also removed those factors that caused one tissue to have an oppositely-signed small effect (absolute Z-score < 3, or P value > 0.00135) when compared to the factor where this eQTL has the strongest effect; such discrepancies may often reflect allelic heterogeneity or LD contamination rather than true opposite effects from the same causal variant [18]

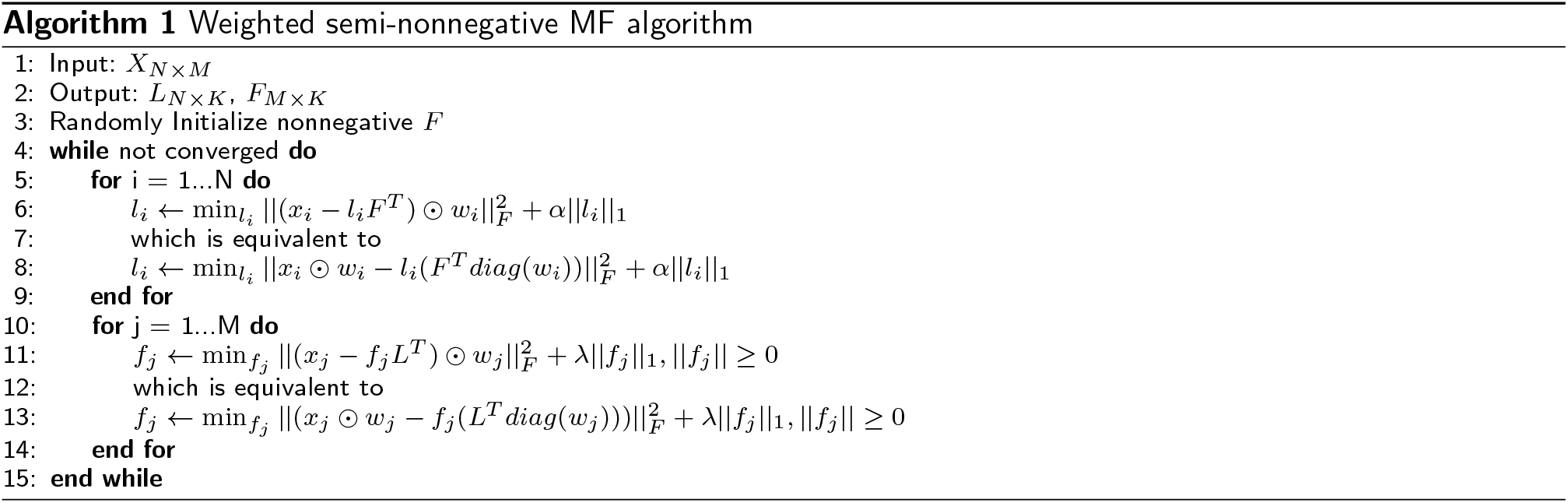

### Background SNP-gene pairs

For enrichment analyses, random SNP-gene pairs were sampled from all SNP-gene pairs to match for eQTLs by three criteria: (1) SNP MAF was matched to the eQTL variants’ MAF, (2) distance from the SNP to TSS of the gene was matched to eQTL, and (3) number of SNPs per gene were matched as in eQTLs.

### Enrichment analysis of eQTLs in different chromatin states

For each 5bp window centered on each SNP, we identified overlapping (1) chromatin state predictions from the Roadmap Epigenomics project and (2) regions of open chromatin identified by DNAse-seq from ENCODE[26, 27, 63, 64, 65]. In Roadmap, chromatin states are predicted for each tissue or cell type that include enhancers, promoters, and transcribed regions. We used the standard 15-state Roadmap segmentations independently for each of the samples that were matched to GTEx tissues (Additional file 2: Table S2, S3). If a tissue had more than one data set available, we merged the data sets using BEDTools [66]. For the data sets using genome assembly hg19, we used liftOver to map the peaks to GRCh38 [67]. We built the 2 × 2 contingency table for eQTLs from each factor and across the 15 chromatin states. In the table, the first row includes eQTL variants in the factor, and the second row includes randomly matched SNPs. The columns indicate number of SNPs that are located in the tested chromatin state in the tested tissues. Both tissues matched for the factor and tissues not matched for the factor were tested. We then ran a two-sided Fisher’s exact test for each contingency table and corrected the P values using BH-correction. To summarize the results across tissues and across factors, we used a random-effects model (`rma()` in R) to obtain the combined odds ratio and combined standard error [68].

### Use thresholding to derive u-eQTLs and ts-eQTLs

We defined ts-eQTLs in *a* tissue as those with P value > 0.001 in at least 44 other tissues, and with P value < 100 × the most extreme P value of the eGene in *the* tissue (Additional file 1: Figure S3). The thresholds were chosen such that we have a reasonable number of ts-eQTLs, and at the same time only eQTLs with a high probability of being casual were included. U-eQTLs were restricted to those found in the credible sets for at least 5 tissues.

### Enrichment analysis of transcription factor binding sites

To examine the enrichment of TF binding sites in u-eQTLs and in ts-eQTLs, we constructed the 2 × 2 contingency tables across factors for each TF. For each TF, we first annotated its binding sites by overlapping tissue-specific enhancer predictions from RoadMap and its TFBS predictions on the genome from JASPAR [25, 26]. We then used genes with at least one variant located in TFBS to avoid genes intrinsically lacking variants in TFBS. In the contingency table for each TF, the first row includes eQTLs, and the second row includes randomly matched SNP gene pairs. For u-eQTLs, the columns indicate the number of genes with or without universal variants in the TFBS. For ts-eQTLs, first column indicates the number of genes with or without tissue-specific variants in the TFBS. One thing to note is that the TFBS were annotated using matched tissues for each factor. Fisher’s exact test was performed for each of these contingency tables, and the P values were corrected using BH-correction.

For eQTLs from each factor, the analysis was done for TFs with median TPM > 1 in at least half of the corresponding tissues with available data. TFs with a total number of genes in TFBS < 10 were removed. We checked to show that the tissue specificity of the enriched TFs is unlikely to result from filtering TFs based on expression level and the number of hits (Additional file 1: Figure S21, S22).

### Identification of allele specific binding sites using ChlP-seq data

FASTQ files from human liver samples of HNF4A and CTCF were downloaded from ENCODE web portal and aligned to GRCh38 genome assembly by STAR [69] (Additional file 2: Table S4). Reads that mapped to variants in GTEx and passed WASP filters were extracted [70]. BAM files of the samples and controls from the same ENCODE repository were downloaded and peak-calling was performed using MACS2 [71]. Only reads that mapped to peaks at q-value < 0.1 were included and ASB was computed for each variant with more than 10 reads by examining if the numbers of reads at each allele were significantly different, using a two-tailed binomial test. Variants with significant ASB events were called at BH corrected P value < 0.05.

## Ethics approval and consent to participate

Please refer to The GTEx Consortium 2019, in submission.

## Consent for publication

Not applicable

## Availability of data and materials

The dataset supporting the conclusions of this article is available to authorized users via dbGaP under accession phs000424.v8 and on the GTEx portal (http://gtexportal.org/). All the code used for the matrix factorization and mapping eQTLs is available on GitHub (https://github.com/heyuan7676/ts_eQTLs).

## Author’s contributions

A.B., C.D.B. and Y.H. designed and coordinated the project. Y.H. performed the analysis. Y.H., A.B. and C.D.B. wrote the manuscript. S.B.C. participated in data analysis and critically revised the manuscript. M.A. participated in the analysis of GWAS hits. K.S. participated in running flashr. F.A. and K.G.A. were responsible for V8 data generation and cis-eQTL calling. A.N.B., R.B. and H.K.I harmonized and imputed GWAS summary statistics. A.B. and C.D.B. conceived and supervised the study. All authors read and approved the final manuscript.

## Acknowledgements

A.B. is supported by NIMH 1R01MH109905, NHGRI 1R01HG010480 and Searle Scholar’s Program. C.D.B. is supported by RF1AG05547701 and R01HL133218. M.A. is supported by T32 HL007227. F.A., K.G.A. are supported by GTEx program grants HHSN268201000029C, 5U41HG009494. H.K.I. is supported by R01MH107666 and P30DK020595. The Genotype-Tissue Expression (GTEx) project was supported by the Common Fund of the Office of the Director of the National Institutes of Health (NIH). Additional funds from the National Cancer Institute; National Human Genome Research Institute (NHGRI); National Heart, Lung, and Blood Institute; National Institute on Drug Abuse; National Institute of Mental Health; and National Institute of Neurological Disorders and Stroke. Donors were enrolled at Biospecimen Source Sites funded by Leidos Biomedical, Inc. (Leidos) subcontracts to the National Disease Research Interchange (10XS170) and Roswell Park Cancer Institute (10XS171). The Laboratory, Data Analysis and Coordinating Center (LDACC) was funded through a contract (HHSN268201000029C) to The Broad Institute, Inc. Biorepository operations were funded through a Leidos subcontract to Van Andel Institute (10ST1035). Additional data repository and project management provided by Leidos (HHSN261200800001E).

## Additional Files

Additional file 1

Supplementary figure. Fig S1 - S22.

Additional file 2

Supplementary tables. Table S1 - S4.

Additional file 3

GTEx Consortium Information.

