## Additional file 1 for "Mechanisms of tissue-specific genetic regulation revealed by latent factors across eQTLs"

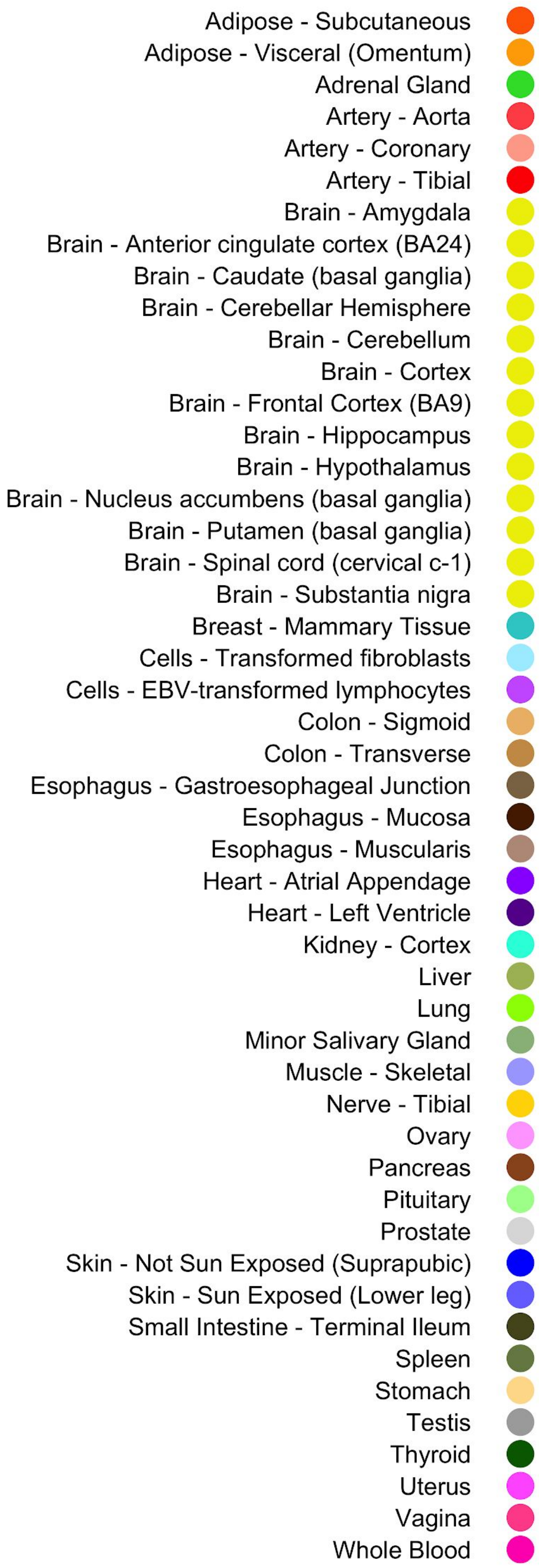


**Figure S1.** Color scheme of 49 tissues in GTEx.

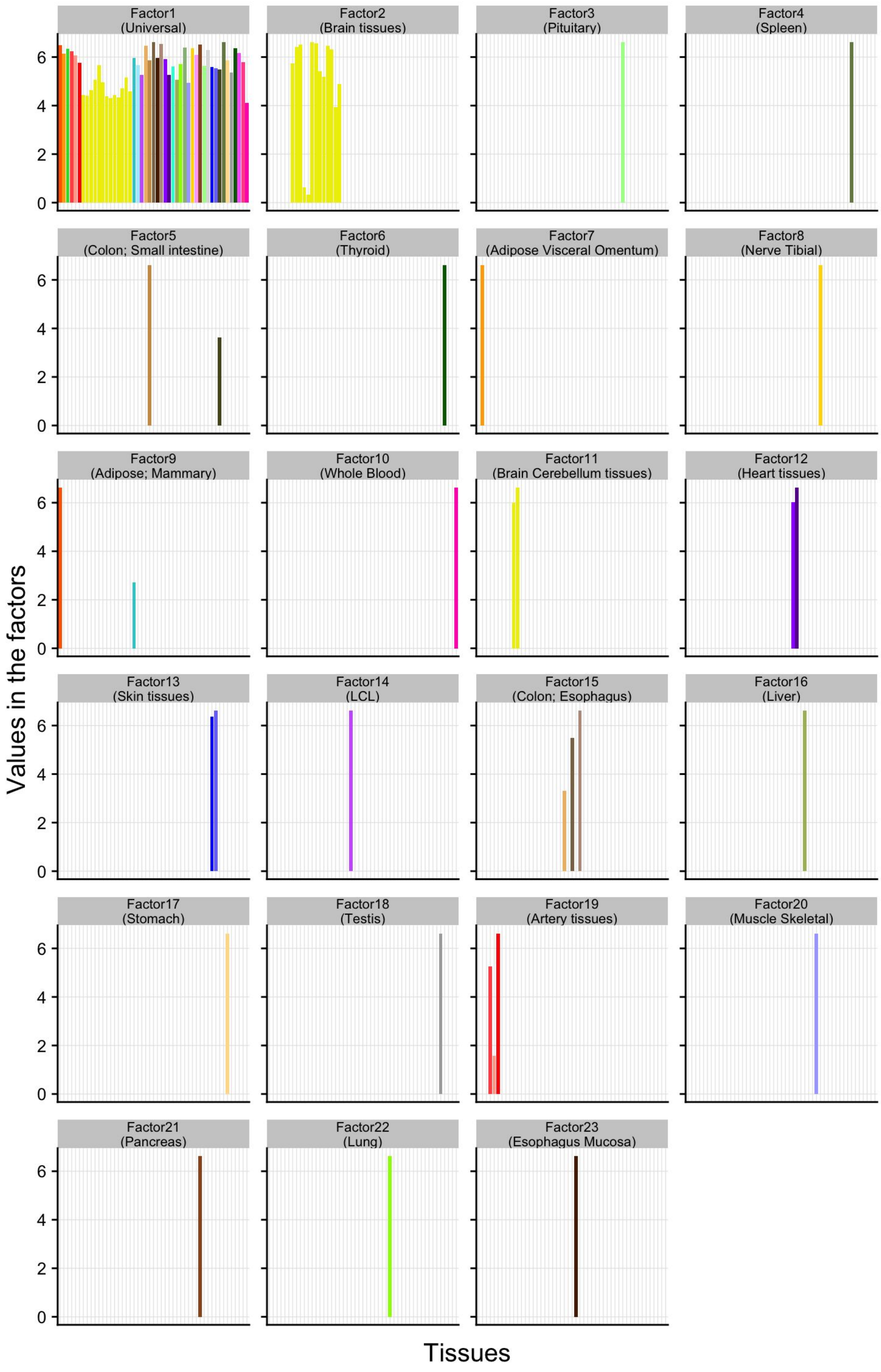


**Figure S2.** Learned factors from sn-spMF. Each panel represents one factor. The factors were named by the tissues with non-zero values in them.

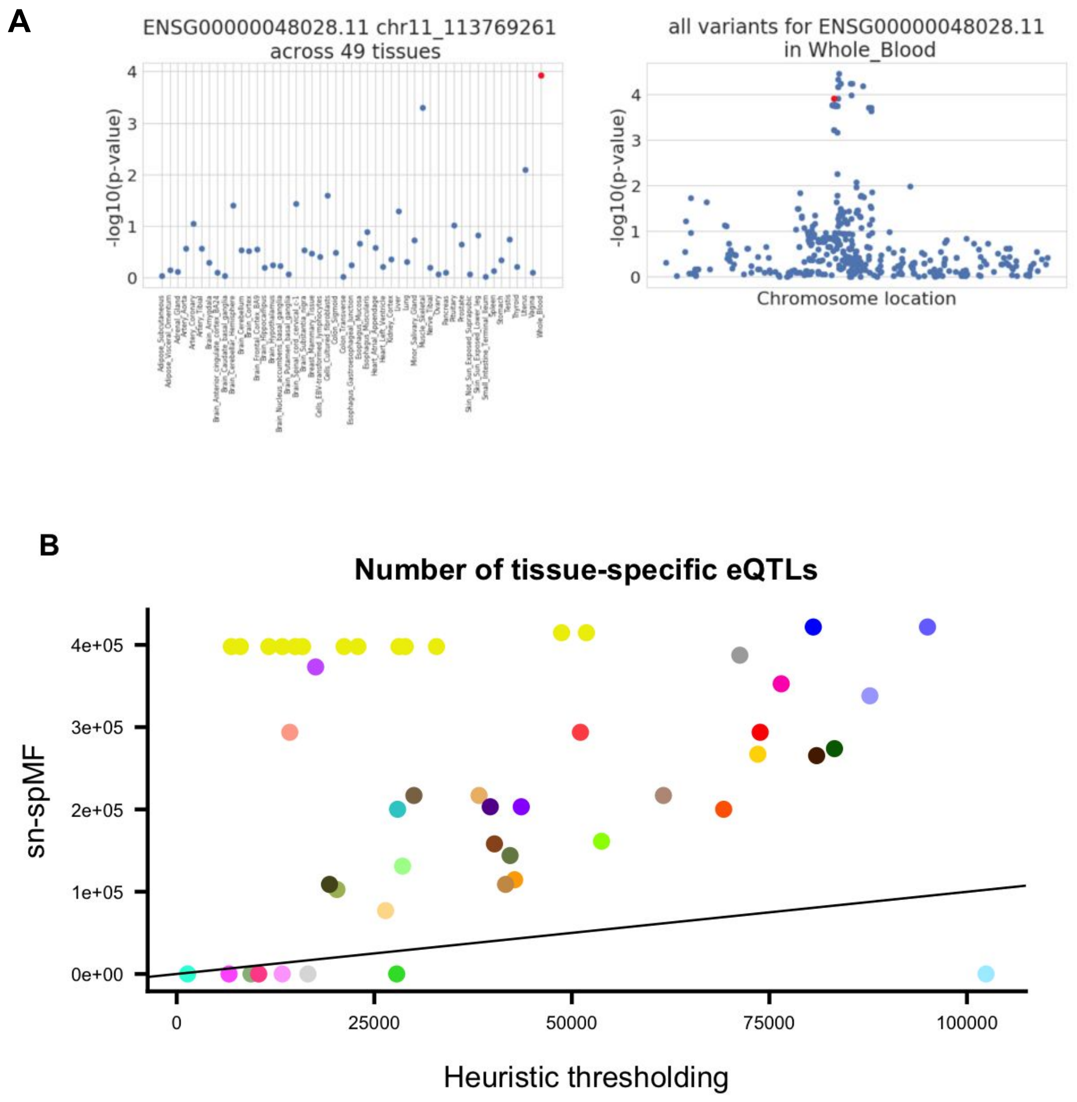


**Figure S3.** U-eQTLs and ts-eQTLs from heuristic methods. **A** An example of whole blood specific eQTL found by thresholding. The red dot represents the found ts-eQTL. Left panel: eQTL results for this SNP gene pair from 49 tissues (such that ts-eQTL signal is presented in few other tissues); Right panel: eQTL results for all variants of the eGene in whole blood (such that the ts-eQTL likely to be casual). **B** Compare the number of ts-eQTLs from the heuristic thresholding method and sn-spMF.

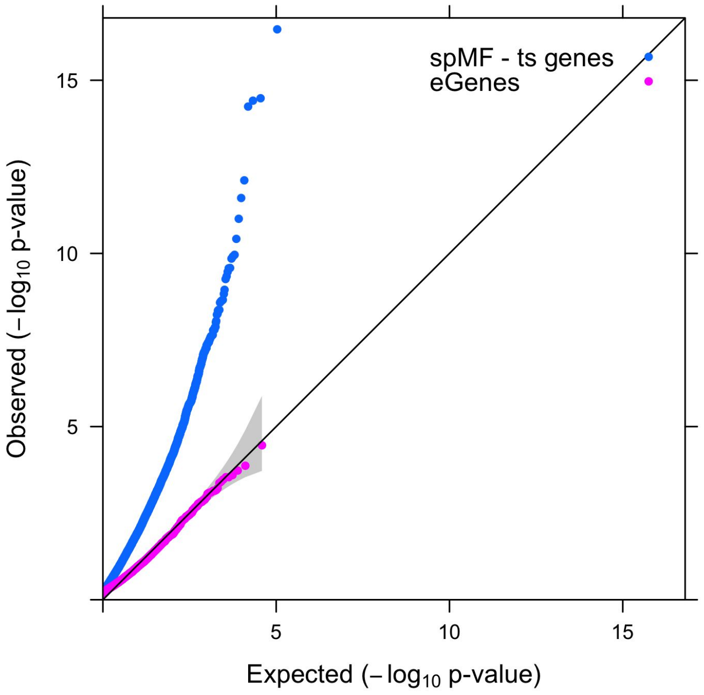


**Figure S4.** QQplot of -log10(p-value) for gene set enrichment analysis (GSEA) for ts-eQTL genes from sn-spMF and eGenes for all the tissues. For each factor from sn-spMF, only the genes that were tested for the corresponding tissues were included as the background. For eGenes, genes tested for eQTL analysis were used as the background. At FDR < 0.05, 546 unique genesets are enriched for tissue-specific genes, while no enrichment is found for the eGenes for each tissue.

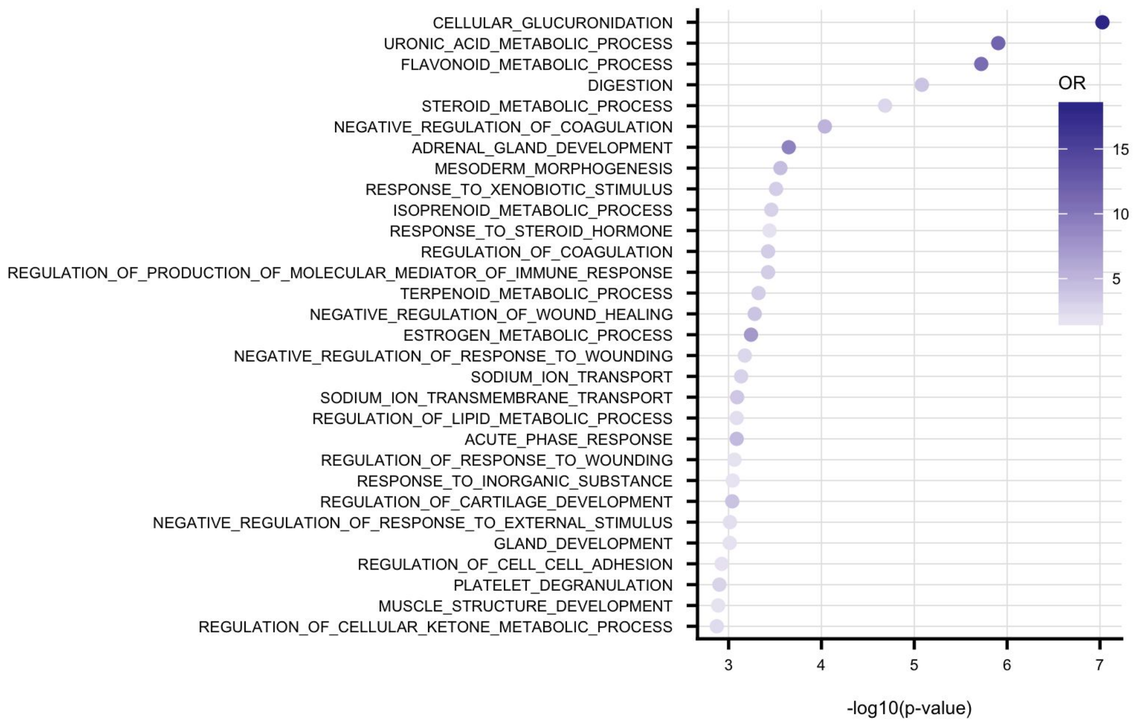


**Figure S5.** Top 30 enriched GO terms for genes with liver specific eQTLs and no u-eQTL.

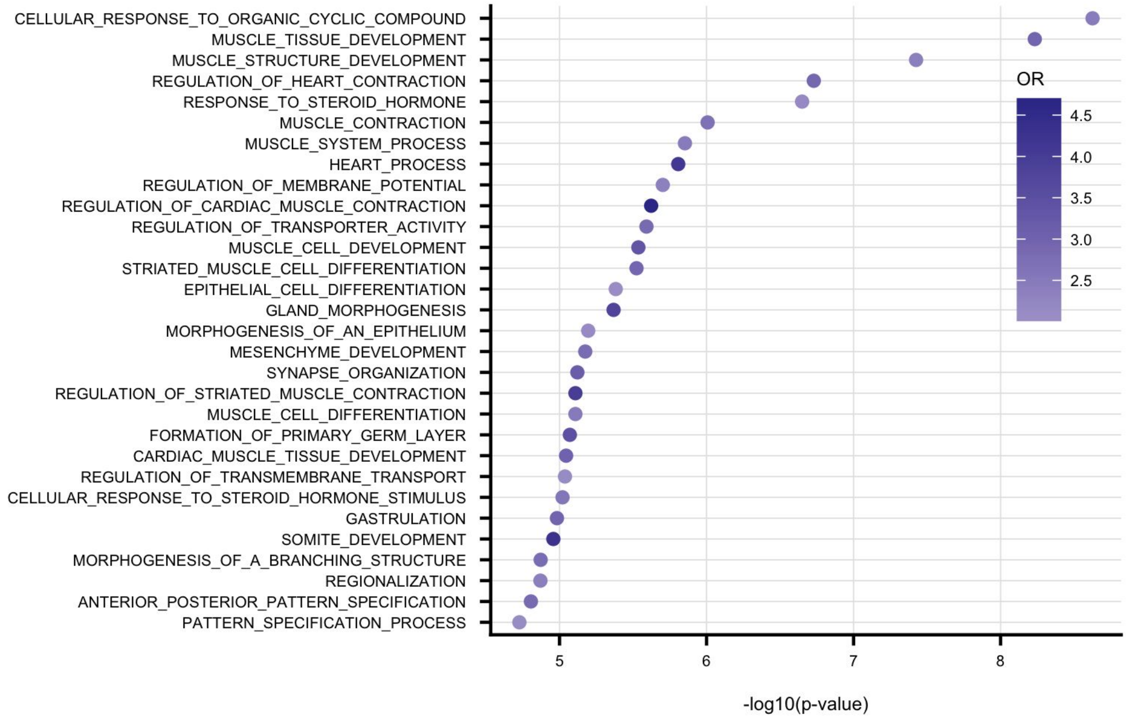


**Figure S6.** Top 30 enriched GO terms for genes with heart specific eQTLs and no u-eQTL.

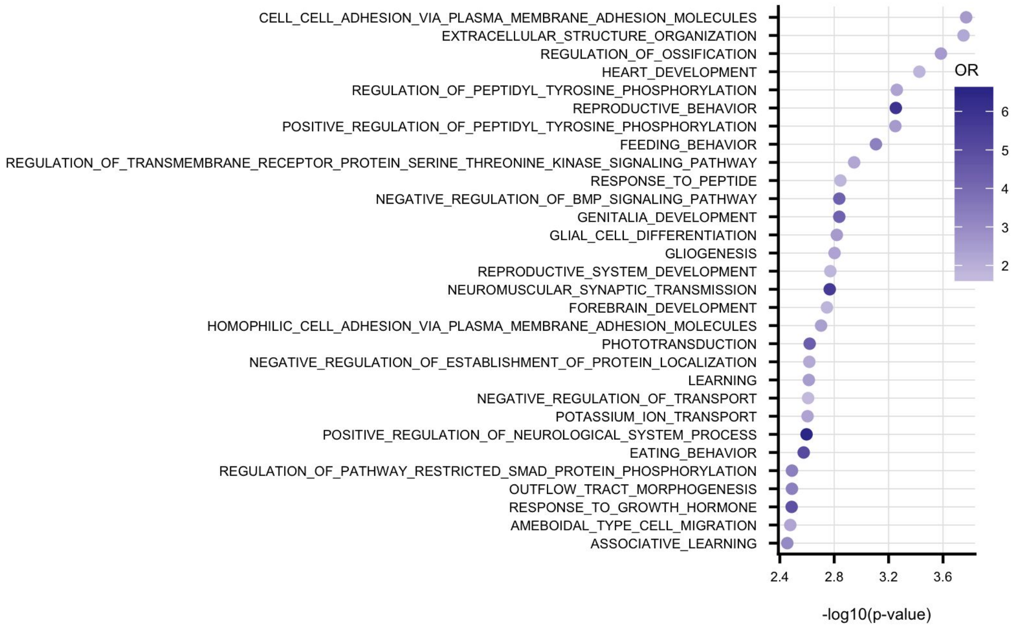


**Figure S7.** Top 30 enriched GO terms for genes with brain tissues specific eQTLs and no u-eQTL.

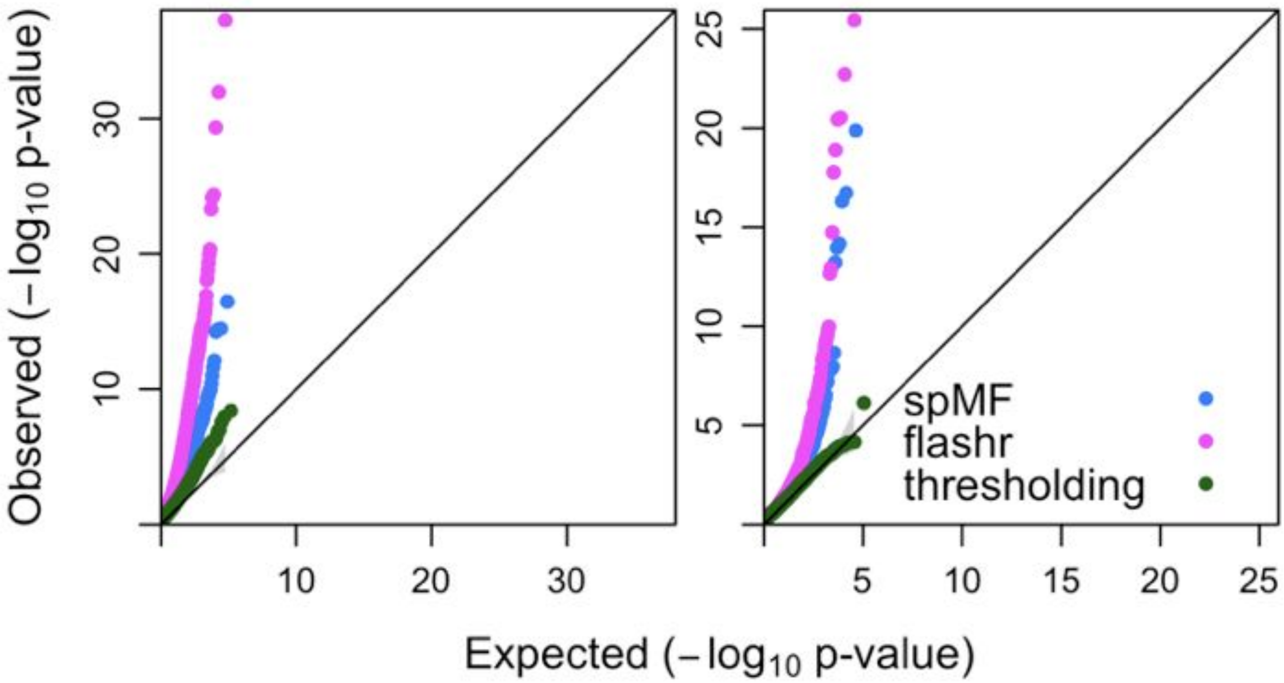


**Figure S8.** QQplot of -log10(p-value) for tissue-specific genes from nn-spMF, flashr and thresholding. **A** Results using tissue-specific genes with tissue-specific eQTLs and with no universal eQTL. **B** Results using tissue-specific genes that appear in less than X factors (X = 6 for sn-spMF and flashr, X = 7 for threshodling for comparable size of the gene sets). At FDR < 0.05, 49 GO terms are enriched for sn-spMF and 85 GO terms are enriched for flashr, and no GO term is enriched for heuristic method.

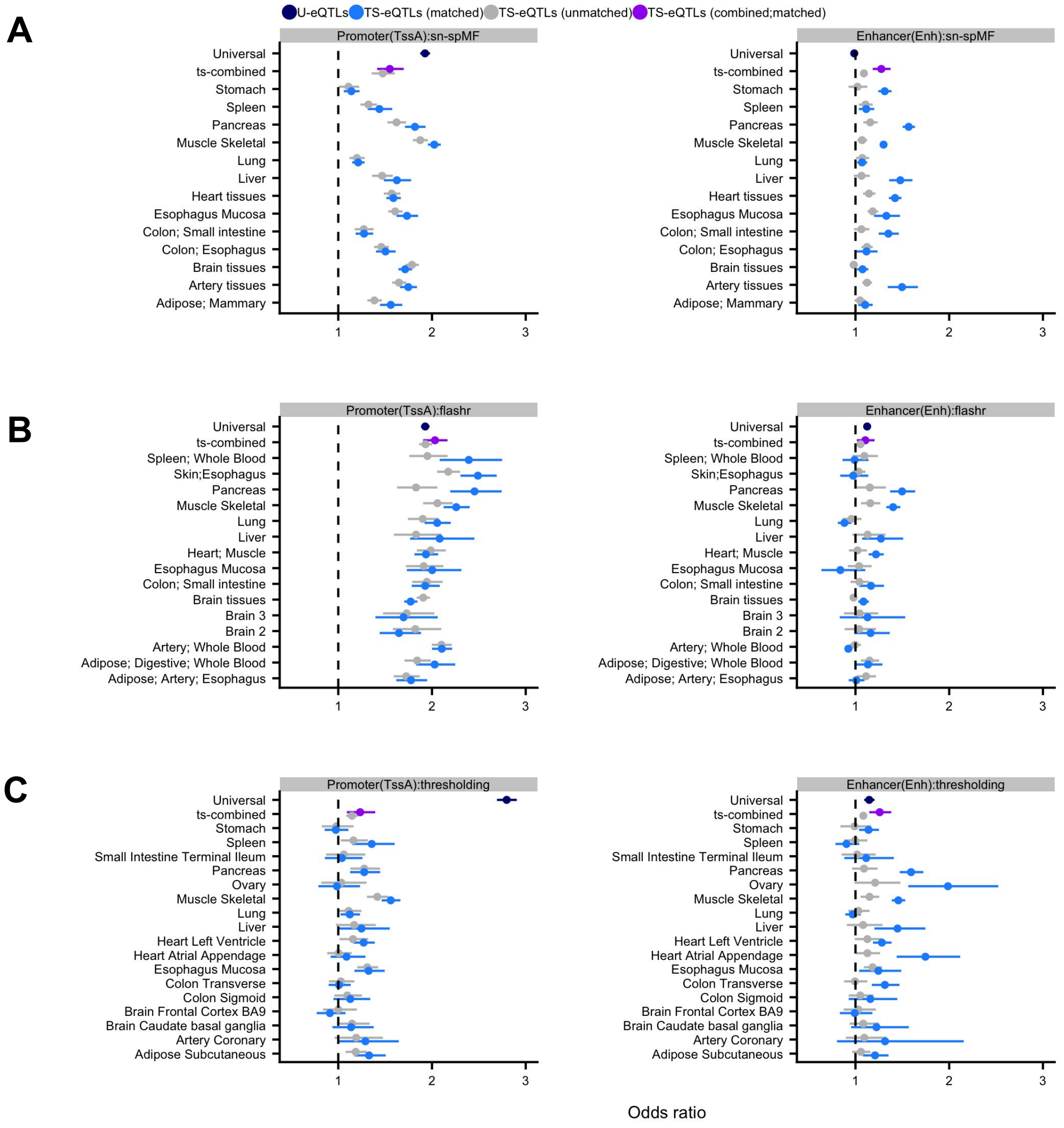


**Figure S9.** Enrichment analysis of chromosome regions for u-eQTL variants and ts-eQTL variants from sn-spMF, flashr and heuristic thresholding. **A** For sn-spMF, ts-eQTL variants are more enriched in enhancers than u-eQTL variants (OR=1.3 and OR=1.0), while u-eQTL variants are more enriched in promoters than ts-eQTL variants (OR=1.9 and OR=1.6) . **B** For flashr, both u-eQTL and ts-eQTL variants are enriched in promoters (OR=1.9 and OR=2.0), but weakly enriched for enhancers, with ts-eQTLs not demonstrating greater enrichment than u-eQTLs (OR=1.1 and OR =1.1. This may be partially due to the somewhat denser factors of flashr not identifying eQTLs that are as highly tissue specific. **C** For heuristic thresholding, ts-eQTL variants are more enriched in enhancers than u-eQTL variants (OR=1.3 and OR=1.1), while u-eQTL variants are more enriched in promoters than ts-eQTL variants (OR=2.8 and OR=1.2) .

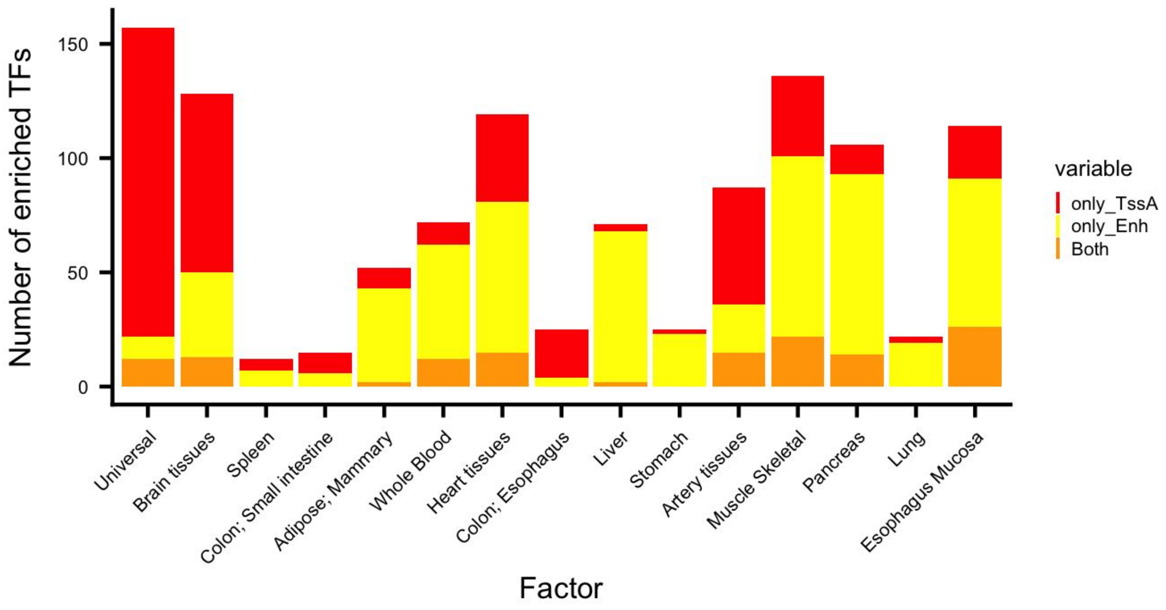


**Figure S10.** Enriched TFs in only promoters, or only in enhancers, or in both promoters and enhancers for eQTLs across factors from sn-spMF.

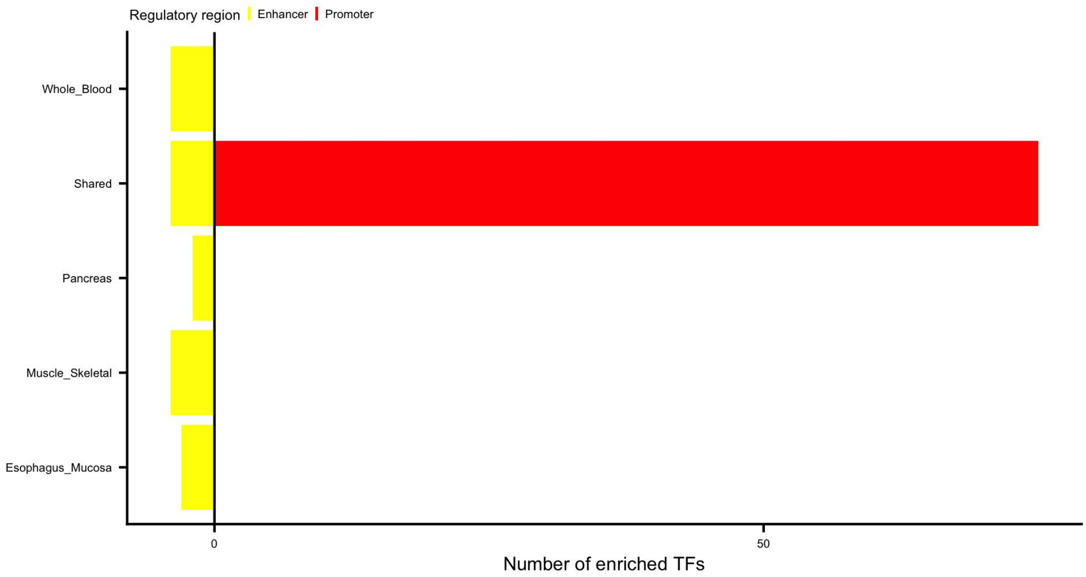


**Figure S11.** Number of enriched TFs in eQTLs from heuristic thresholding in enhancers and promoters.

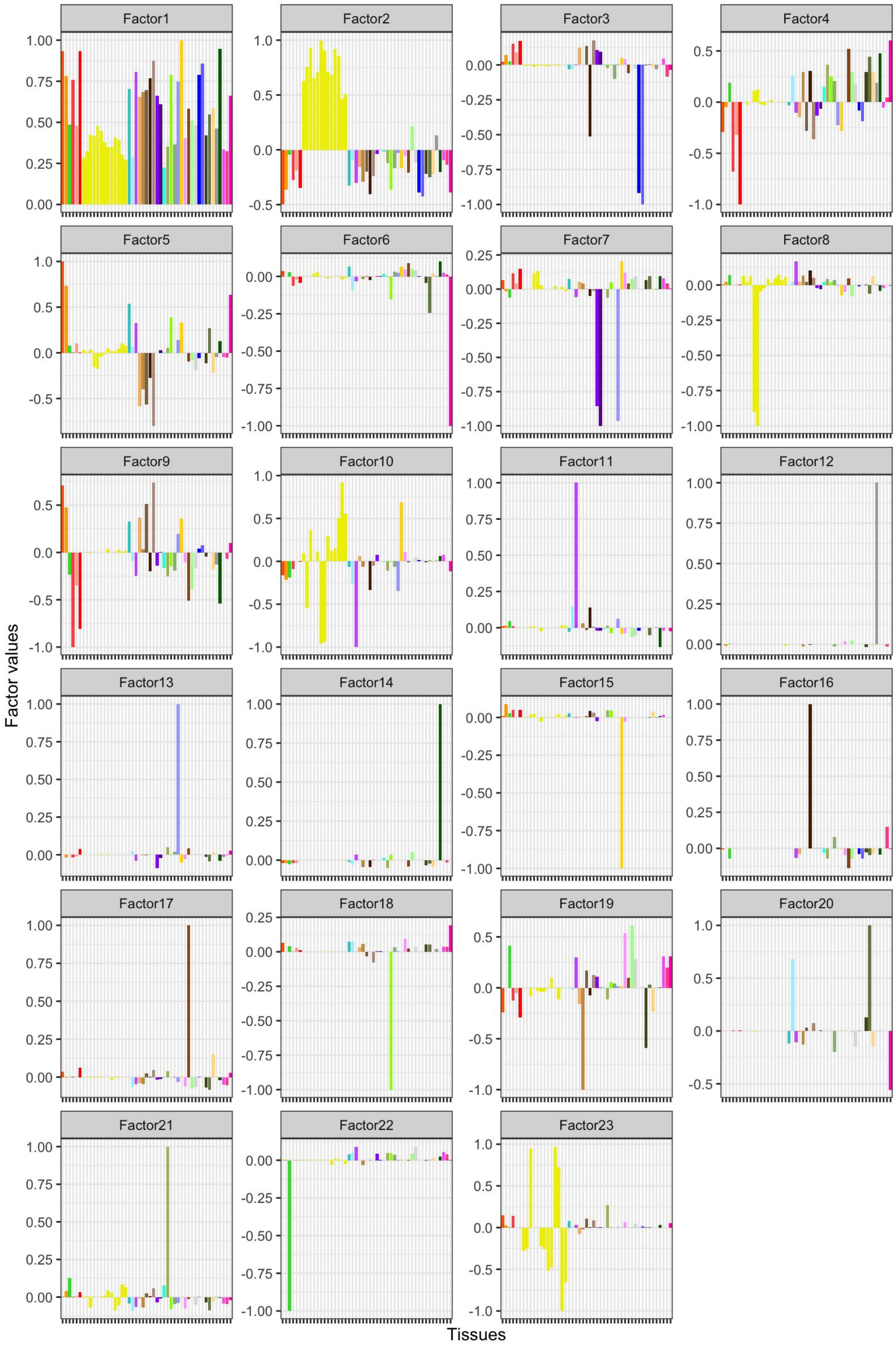


**Figure S12.** Factor matrix learned from flashr using the same input eQTLs as sn-spMF.

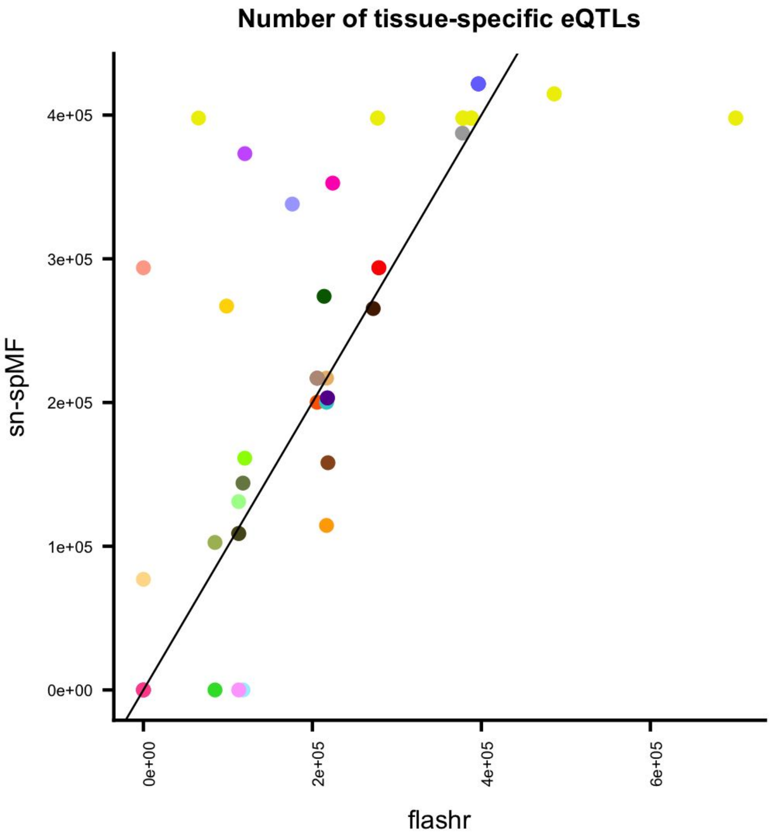


**Figure S13.** Compare the number of ts-eQTLs from flashr and sn-spMF.

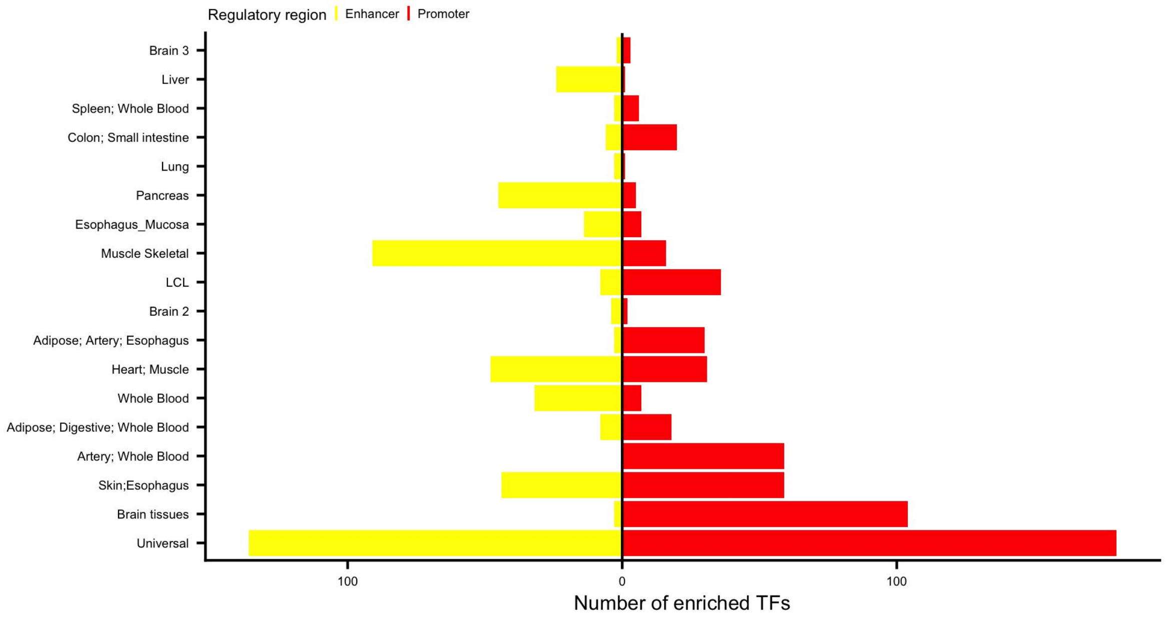


**Figure S14.** Number of enriched TFs in eQTLs from flashr in enhancers and promoters.

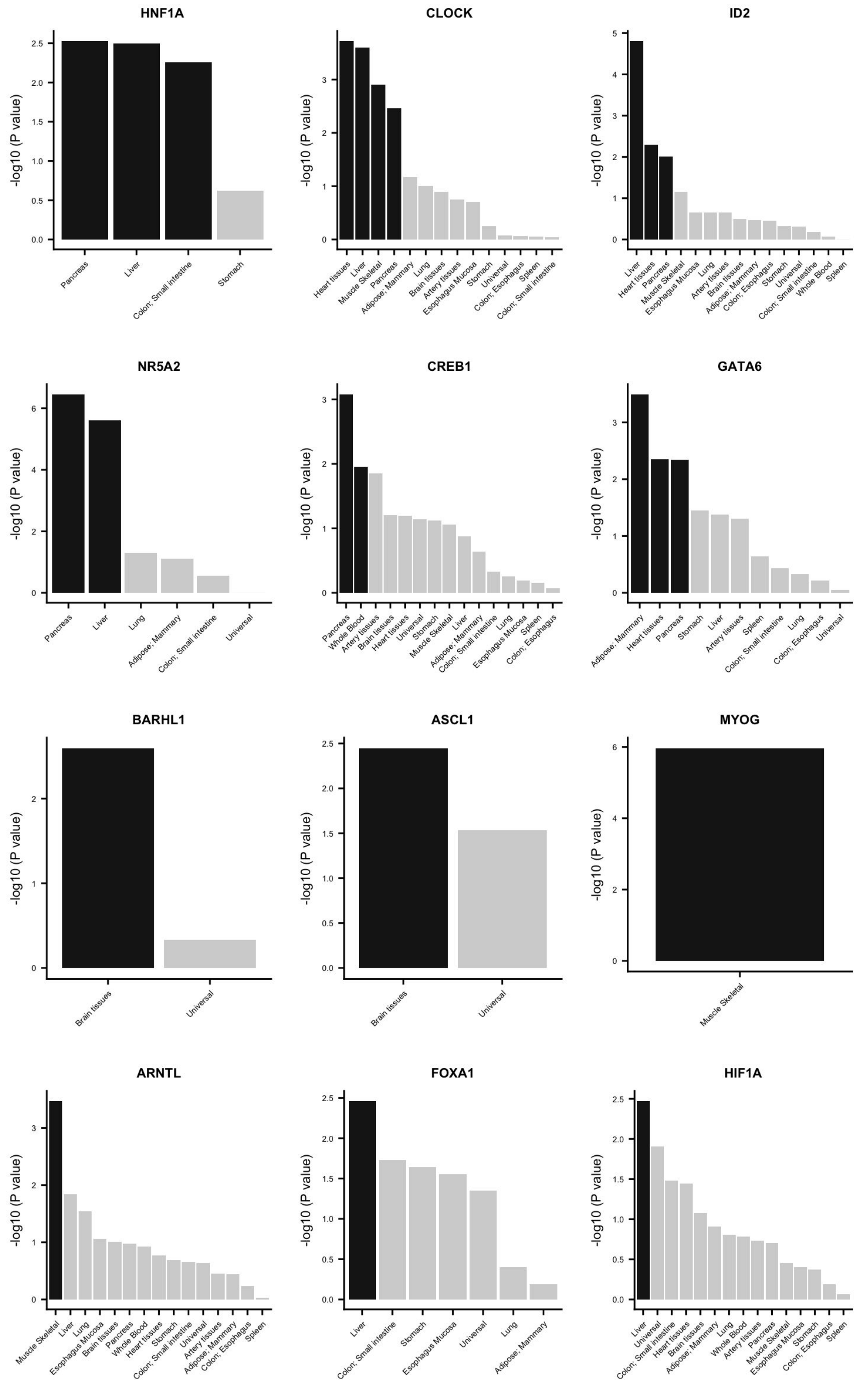


**Figure S15.** More examples of TFs with known functions that are enriched in ts-eQTLs of the related tissues

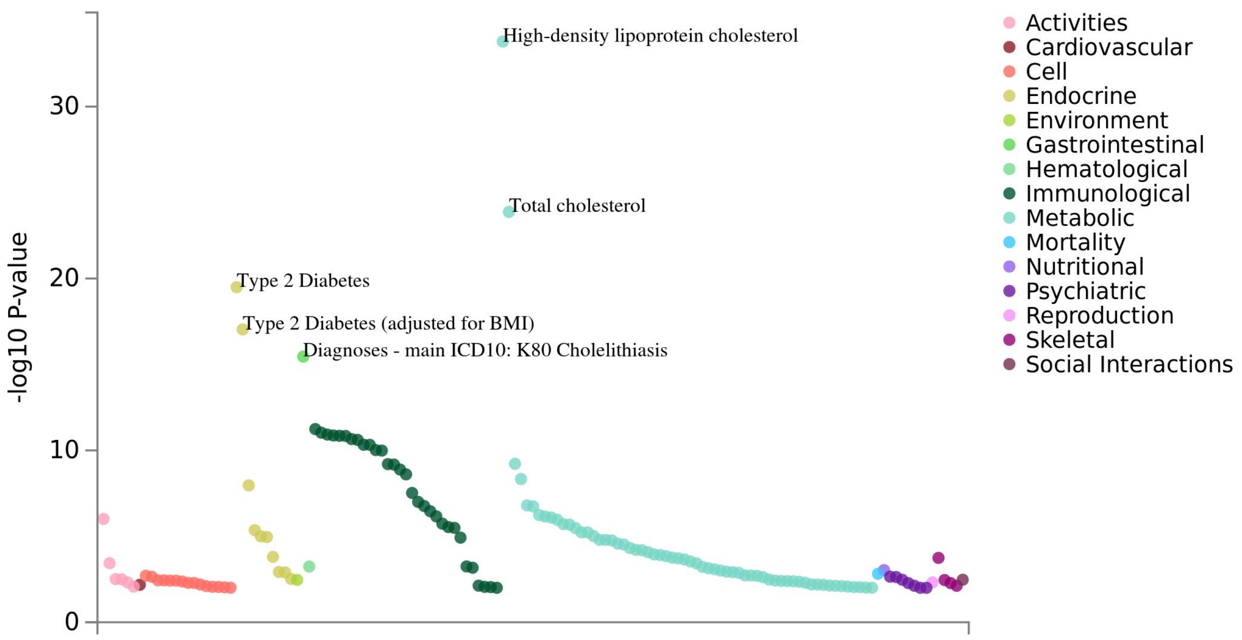


**Figure S16.** PheWAS plot for the missense variant rs1800961 in HNF4A.

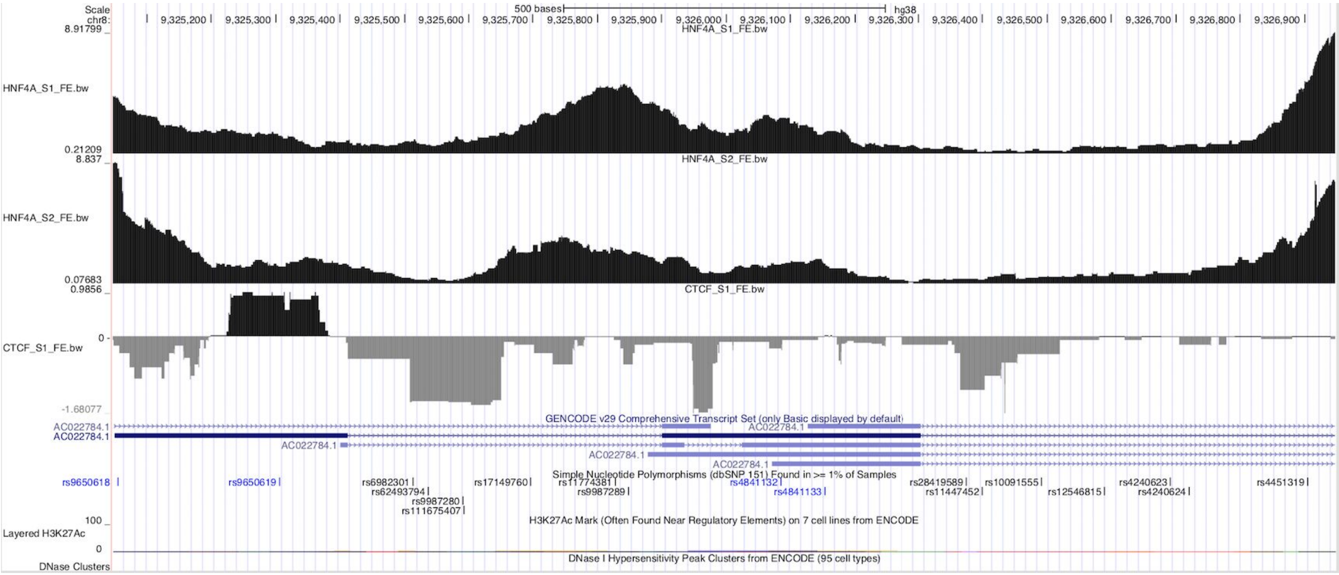


**Figure S17.** ChIP-seq data of HNF4A and CTCF shows that rs9987289 is in the peak region of HNF4A, but not in the peak region of CTCF.

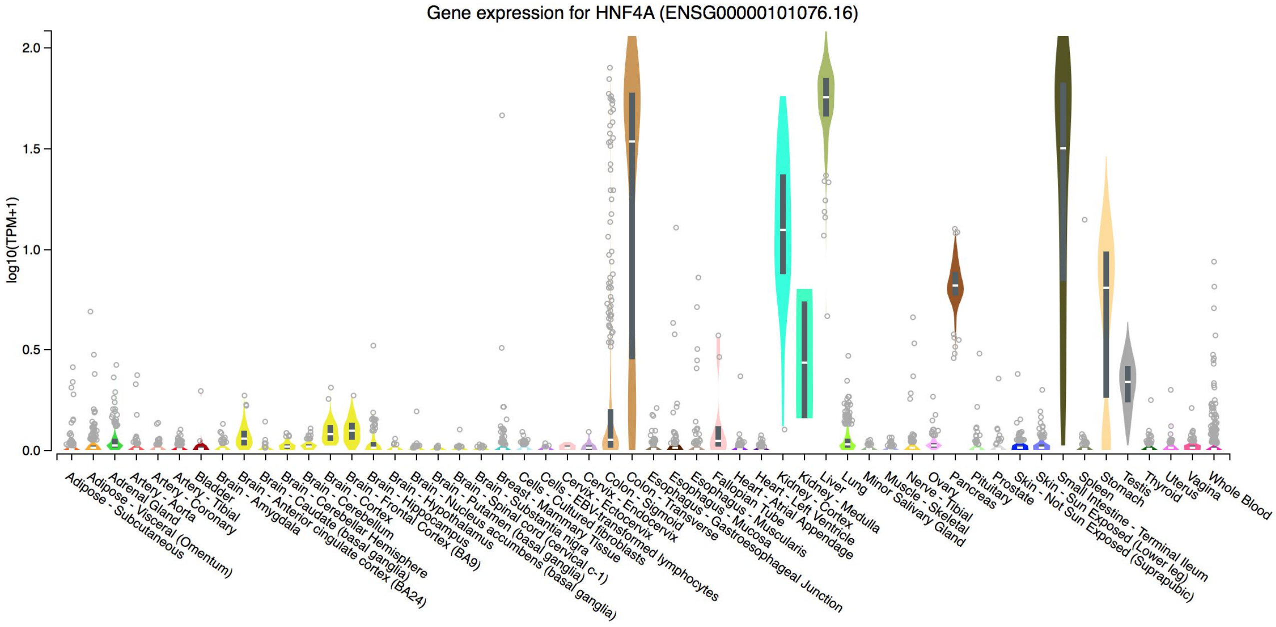


**Figure S18.** Expression level of HNF4A across tissues from GTEx v8.

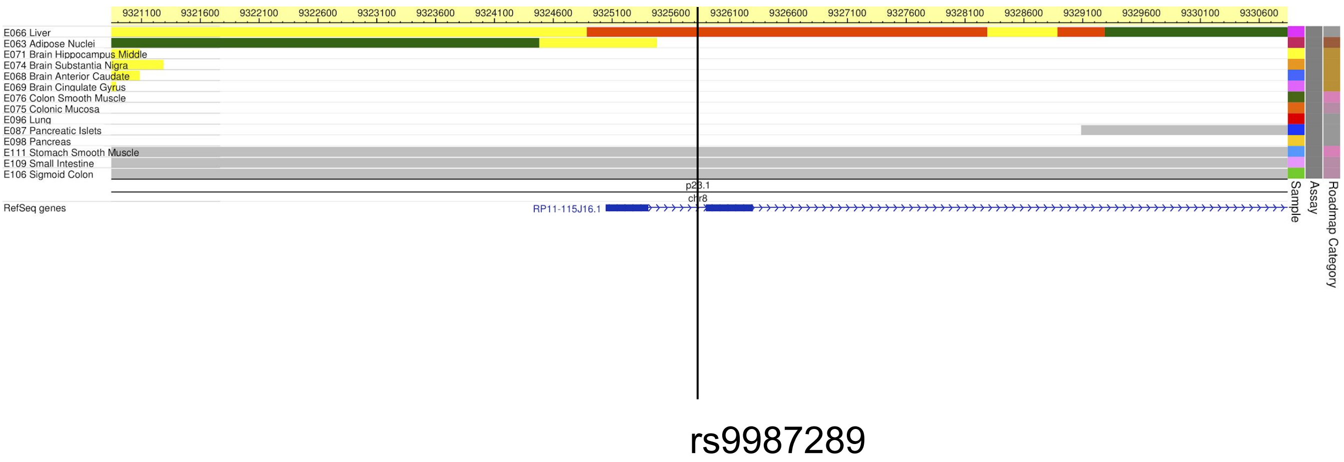


**Figure S19.** Prediction of chromatin states from Roadmap Project.

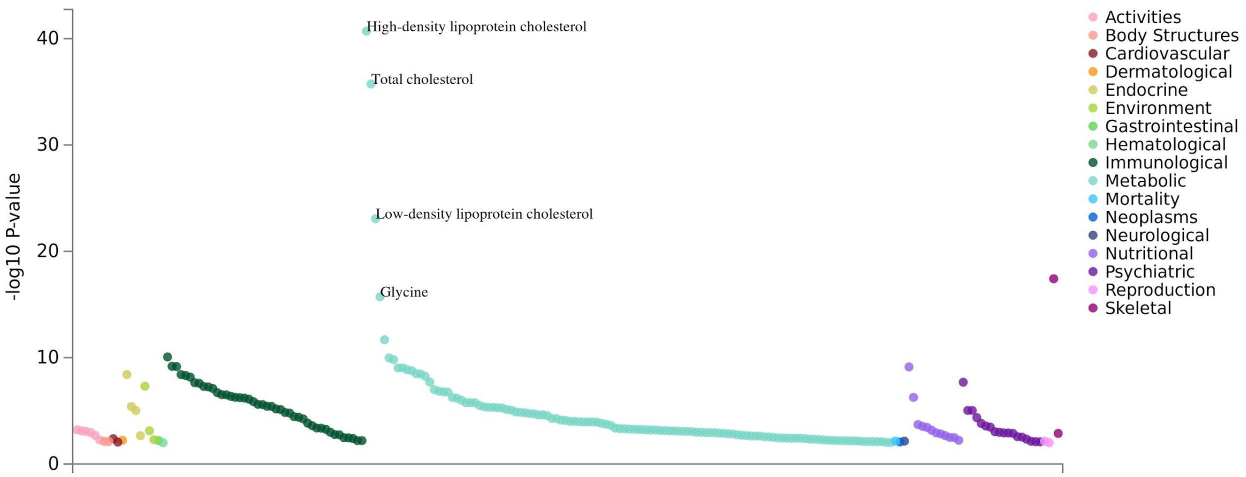


**Figure S20.** PheWAS plot for the liver-specific variant rs9987289.

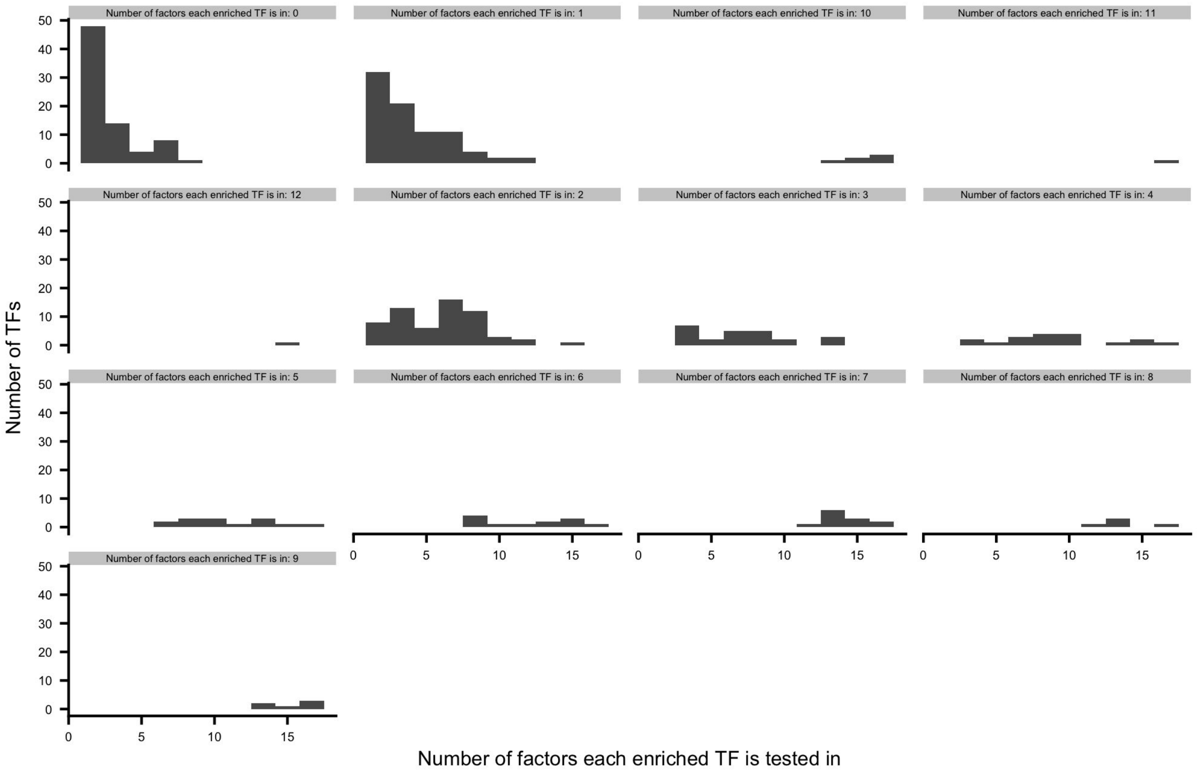


**Figure S21.** Number of factors each each TF is enriched in in promoter, and the number of factors each TF is tested in.

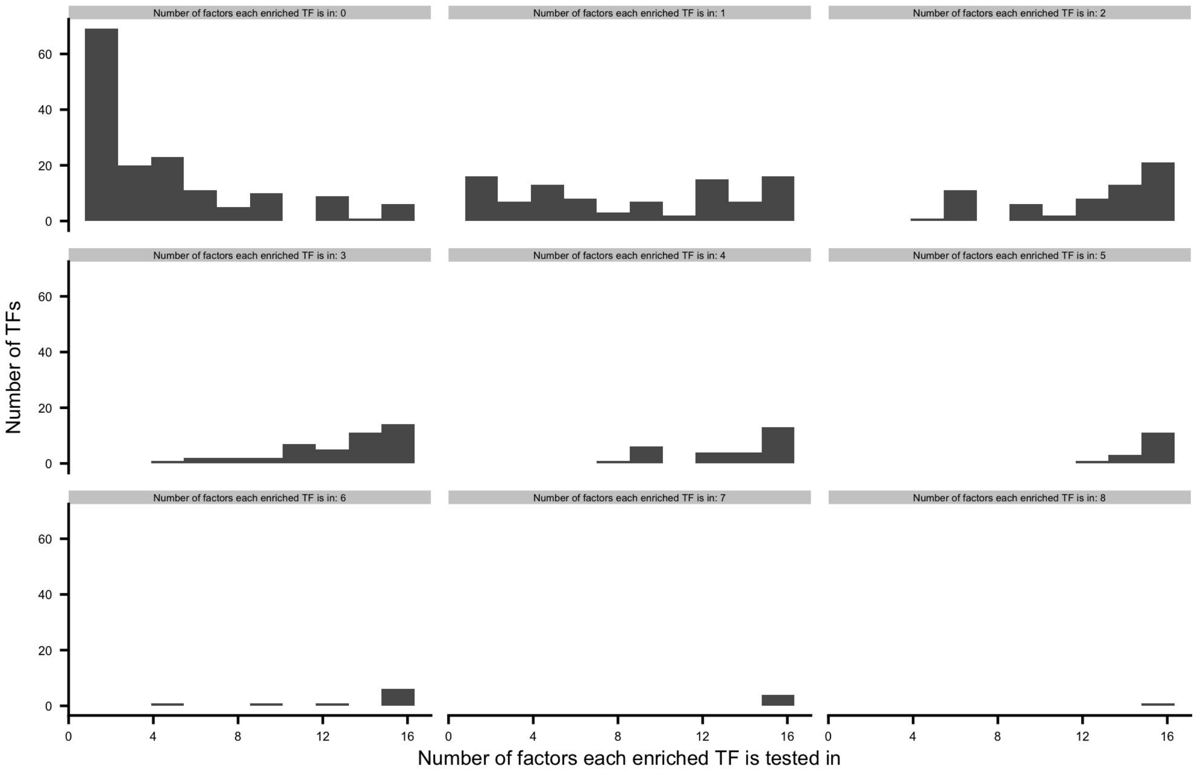


**Figure S22.** Number of factors each each TF is enriched in in enhancer, and the number of factors each TF is tested in.
