## Additional file 2 for "Mechanisms of tissue-specific genetic regulation revealed by latent factors across eQTLs"

Supplementary Table 1. Enriched TFs with strong literature support

| TF | Tissues | Reference (DOI) |
|---|---|---|
| BCL6 | Adipose; Mammary | 10.1073/pnas.1907308116 |
| SREBF1 | Adipose; Mammary | 10.1038/srep00178 |
| TWIST1 | Adipose; Mammary | 10.1016/j.cell.2009.01.051 |
| SMAD4 | Artery tissues | 10.1128/MCB.00577-07 |
| TWIST1 | Artery tissues | 10.1161/CIRCRESAHA.116.308870 |
| LHX2 | Brain tissues | 10.1523/JNEUROSCI.3145-15.2016 |
| OLIG1 | Brain tissues | 10.1038/nn.2600 |
| SOX1 | Brain tissues | 10.1016/s0306-4522(03)00158-1 |
| SOX2 | Brain tissues | 10.1242/dev.01204 |
| SOX6 | Brain tissues | 10.1038/nn.2387 |
| SOX9 | Brain tissues | 10.1523/JNEUROSCI.3199-16.2017 |
| XBP1 | Brain tissues | 10.2119/molmed.2016.00229. |
| BCL6 | Heart tissues | 10.1016/S0008-6363(99)00007-3 |
| CLOCK | Heart tissues | 10.1161/hh1101.091190 |
| FOXO1 | Heart tissues | 10.1093/cvr/cvs426 |
| FOXP1 | Heart tissues | 10.1002/humu.22366 |
| ID2 | Heart tissues | 10.1016/j.cell.2007.04.036 |
| SOX6 | Heart tissues | 10.1073/pnas.97.8.4180 |
| SOX9 | Heart tissues | 10.1073/pnas.0401711101 |
| TWIST1 | Heart tissues | 10.1016/j.ydbio.2010.08.021 |
| XBP1 | Heart tissues | 10.1111/acel.12460 |
| CLOCK | Liver | 10.1074/jbc.M304564200 |
| FOXA1 | Liver | 10.1038/nature03649 |
| FOXO1 | Liver | 10.1038/nm.2049 |
| ID2 | Liver | 10.1074/jbc.M109.013961 |
| MAFG | Liver | 10.1016/j.cmet.2015.01.007 |
| NFIL3 | Liver | 10.1016/j.metabol.2017.08.007 |
| NR5A2 | Liver | 10.1016/j.ydbio.2016.07.019 |
| XBP1 | Liver | 10.1074/jbc.M115.676239 |
| CLOCK | Muscle Skeletal | 10.1073/pnas.1014523107 |

| | | |
|---|---|---|
| FOXO1 | Muscle Skeletal | 10.18632/oncotarget.12891 |
| PITX1 | Muscle Skeletal | 10.1016/j.ydbio.2006.06.055 |
| SOX6 | Muscle Skeletal | 10.1002/dvdy.21223 |
| SREBF1 | Muscle Skeletal | 10.1371/journal.pone.0050878 |
| TEAD1 | Muscle Skeletal | 10.1074/jbc.M113.515817 |
| ATF4 | Pancreas | 10.1016/j.cmet.2008.01.008 |
| CLOCK | Pancreas | 10.1038/nature09253 |
| FOXO1 | Pancreas | 10.1210/en.2015-1852 |
| FOXP1 | Pancreas | 10.1007/s00125-015-3635-3 |
| ID2 | Pancreas | 10.1007/s12020-008-9039-0 |
| NEUROD1 | Pancreas | 10.1101/gad.9.8.1009 |
| NKX6-1 | Pancreas | 10.1016/j.celrep.2013.08.010 |
| NR5A2 | Pancreas | 10.1016/j.ydbio.2016.07.019 |
| SOX6 | Pancreas | 10.1074/jbc.M700460200 |
| SREBF1 | Pancreas | 10.1194/jlr.M700533-JLR200 |
| TEAD1 | Pancreas | 10.1038/ncb3160 |
| XBP1 | Pancreas | 10.1038/sj.emboj.7600903 |
| FLI1 | Whole Blood | 10.1016/j.cub.2008.07.048 |
| FOXO1 | Whole Blood | 10.1038/ncomms11023 |
| NFIL3 | Whole Blood | 10.1136/annrheumdis-2018-213764 |
| RUNX1 | Whole Blood | 10.1038/emboj.2012.275 |

Supplementary Table 2. Samples from Roadmap Epigenomics project mapped to GTEx tissues

| Roadmap EID | Epigenome name | Tissues in GTEx |
|---|---|---|
| E107 | Skeletal muscle male | Muscle Skeletal |
| E108 | Skeletal muscle female | Muscle Skeletal |
| E095 | Left ventricle | Heart Left Ventricle, Heart Atrial Appendate |
| E105 | Right ventricle | Heart Left Ventricle, Heart Atrial Appendate |
| E097 | Ovary | Ovary |
| E066 | Liver | Liver |
| E098 | Pancreas | Pancreas |
| E096 | Lung | Lung |
| E113 | Spleen | Spleen |
| E071 | Brain hippocampus middle | Brain Hippocampus |
| E068 | Brain anterior caudate | Brain Caudate basal ganglia |
| E074 | Brain substantia nigra | Brain Substantia nigra |
| E063 | Adipose nuclei | Adipose Subcutaneous |
| E065 | Aorta | Artery Coronary |
| E079 | Oesophagus | Esophagus Mucosa, Esophagus Gastroesophageal Junction, Esophagus Muscularis |
| E111 | Stomach smooth muscle | Stomach |
| E076 | Colon smooth muscle | Colon Transverse, Colon Sigmoid |
| E106 | Sigmoid colon | Colon Sigmoid |
| E109 | Small intestine | Small Intestine Terminal Ileum |
| E116 | GM12878 lymphoblastoid | Cells EBV-transformed lymphocytes |

Supplementary Table 3. DNase-seq data from ENCODE project

| ENCODE Accession | ENCODE sample tissue | GTEx tissue |
|---|---|---|
| ENCFF958GWR, ENCFF954PTR | Omental fat pad | Adipose Subcutaneous |
| ENCFF042VKK, ENCFF085NOG, ENCFF108XQG, ENCFF217MXO, ENCFF315CSH, ENCFF367BEU, ENCFF587SIS, ENCFF675UKK, ENCFF688ZWO, ENCFF896DOA, ENCFF977OWF | Adrenal gland | Adrenal Gland |
| ENCFF968IAI | Ascending aorta | Artery Aorta |
| ENCFF822UQG, ENCFF178BNR | Coronary artery | Artery Coronary |
| ENCFF048ZGK, ENCFF267DGC | Tibial artery | Artery Tibial |
| ENCFF240ECT | Caudate nucleus | Brain Caudate basal ganglia |
| ENCFF053XFC, ENCFF337NAS | Cerebellar cortex | Brain Cerebellar Hemisphere |
| ENCFF732MQW, ENCFF966DRW | Cerebellum | Brain Cerebellum |
| ENCFF255NTQ, ENCFF611EHQ, ENCFF631HBT, ENCFF855HES | Frontal cortex | Brain Frontal Cortex BA9 |
| ENCFF026XWM | Putamen | Brain Putamen basal ganglia |
| ENCFF421NEH, ENCFF469JHU | Sigmoid colon | Colon Sigmoid |
| ENCFF134KRY, ENCFF159SOA, ENCFF161NFM, ENCFF384WXP, ENCFF791HOY, | Transverse colon | Colon Transverse |
| ENCFF146AEB | Esophagus muscularis mucosa | Esophagus Gastroesophageal Junction, Esophagus Mucosa, Esophagus Muscularis |
| ENCFF146VYU, ENCFF778BRJ, ENCFF794SOC, ENCFF855YGO | Heart left ventricle | Heart Atrial Appendage, Heart Left Ventricle |
| ENCFF172XNI | Left cardiac atrium | |
| ENCFF207NXB, ENCFF289IMF | Heart right ventricle | |

| | | |
|---|---|---|
| ENCFF036JUB, ENCFF153WQN, ENCFF183AEI, ENCFF262FHU, ENCFF270GNM, ENCFF305WVB, ENCFF402THI, ENCFF484XHW, ENCFF578BEO, ENCFF765BJR, ENCFF812GJU, ENCFF845VOI, ENCFF871EKB, ENCFF916NTG, ENCFF932ATD | Kidney | Kidney Cortex |
| ENCFF081JVT, ENCFF468NND, ENCFF512TSJ | Liver | Liver |
| ENCFF475HWF | Right lobe of liver | |
| ENCFF318TOW, ENCFF353SVP, ENCFF439ZRL, ENCFF484YOE, ENCFF588WQL, ENCFF601TZC, ENCFF642HTL, ENCFF671CWO, ENCFF676GRC, ENCFF690UKD, ENCFF796EIB, ENCFF929FIK, ENCFF944FSO | Left lung | Lung |
| ENCFF449XXS, ENCFF486CWL, ENCFF791OUO | Upper lobe of left lung | |
| ENCFF024SOP, ENCFF115HTH, ENCFF148PHO, ENCFF348CJE, ENCFF363XQF, ENCFF395KUT, ENCFF422YFH, ENCFF679QGU, ENCFF811RTH, ENCFF889NTH, ENCFF909JGU, ENCFF913NRZ, ENCFF962JWU, ENCFF978OUM, ENCFF992VNB | Lung | |
| ENCFF157KGS, ENCFF277WMS, ENCFF281HKU, ENCFF352RNR, ENCFF516FXF, ENCFF586UYY, ENCFF604AQG, ENCFF628MPB, ENCFF785ORF, ENCFF796WDQ, ENCFF941EJJ | Right lung | |
| ENCFF028CVN, ENCFF036PYG, ENCFF040WPR, ENCFF041NEG, ENCFF213IAV, ENCFF229KSL, ENCFF246TUN, ENCFF262NZB, ENCFF308QRZ, ENCFF334ENU, ENCFF349CIP, ENCFF475YRW, ENCFF615LEO, ENCFF755PMB, ENCFF771EKC, ENCFF886DDL, ENCFF994YDK | Muscle of arm | Muscle Skeletal |
| ENCFF016PCJ, ENCFF062LJL, ENCFF182YXK, ENCFF191MBC, | Muscle of back | |

| | | |
|---|---|---|
| ENCFF365RKF, ENCFF376WVL, ENCFF417IJL, ENCFF433VTN, ENCFF468TOZ, ENCFF735DNU, ENCFF758REB, ENCFF766PCO, ENCFF831RQP, ENCFF897TZA, ENCFF994ALS | | |
| ENCFF011UBS, ENCFF031RMC, ENCFF058UNN, ENCFF067VHZ, ENCFF138LZU, ENCFF175BCP, ENCFF241QSX, ENCFF283UJD, ENCFF470XXM, ENCFF699XQF, ENCFF874GGX, ENCFF896ZUQ, ENCFF970QZI | Muscle of leg | |
| ENCFF022UVJ, ENCFF383OJO, ENCFF443ZJX | Muscle of trunk | |
| ENCFF614SOO, ENCFF643WVI | Tibial nerve | Nerve Tibial |
| ENCFF111IXG, ENCFF342PZX, ENCFF701ZFB, ENCFF936ENC | Ovary | Ovary |
| ENCFF398ENA, ENCFF535LEW | Pancreas | Pancreas |
| ENCFF569WWH, ENCFF627DYO, ENCFF779JBV, ENCFF963BGI | Body of pancreas | |
| ENCFF228ZTQ | Lower leg skin | Skin Not Sun Exposed Suprapubic,<br>Skin Sun Exposed Lower leg |
| ENCFF019PSW, ENCFF087XDG, ENCFF130CZB, ENCFF274NTF, ENCFF333MTL, ENCFF412ATV, ENCFF424PWV, ENCFF571UWP, ENCFF617TGM, ENCFF720DUQ, ENCFF731WZI, ENCFF758VXS, ENCFF885IBS | Small intestine | Small Intestine Terminal Ileum |
| ENCFF376YIY, ENCFF534XLO | Spleen | Spleen |
| ENCFF009YJE, ENCFF024HZS, ENCFF187BOF, ENCFF227HYU, ENCFF272WHU, ENCFF278ROU, ENCFF376EBF, ENCFF457SNJ, ENCFF523INF, ENCFF556WLI, ENCFF631XSP, ENCFF694LGV, ENCFF709TJW, ENCFF716YVE, ENCFF749DUT, ENCFF751PUA, ENCFF765AZQ, ENCFF785AIA, | Stomach | Stomach |

| | | |
|---|---|---|
| ENCFF885VHR, ENCFF967LJZ, ENCFF988ZPF | | |
| ENCFF018TWY, ENCFF102TYW, ENCFF618EIJ | Testis | Testis |
| ENCFF440OAH, ENCFF460WME, ENCFF652DKF, ENCFF856OWH | Thyroid gland | Thyroid |
| ENCFF514GYQ | Uterus | Uterus |
| ENCFF018IDK, ENCFF329QLI | Vagina | Vagina |

Supplementary Table 4. ChIP-seq data from ENCODE

| TF | Experiment |
|---|---|
| HNF4A | ENCSR445QRF, ENCSR601OGE |
| CTCF | ENCSR254YRM |
