## Additional file 3 for "Mechanisms of tissue-specific genetic regulation revealed by latent factors across eQTLs"

**GTEx Consortium Information**

**Laboratory and Data Analysis Coordinating Center (LDACC):** François Aguet[1], Shankara Anand[1], Kristin G Ardlie[1], Stacey Gabriel[1], Gad Getz[1,2], Aaron Graubert[1], Kane Hadley[1], Robert E Handsaker[3,4,5], Katherine H Huang[1], Seva Kashin[3.4.5], Xiao Li[1], Daniel G MacArthur[4.6], Samuel R Meier[1], Jared L Nedzel[1], Duyen Y Nguyen[1], Ayellet V Segrè[1,7], Ellen Todres[1]
**Analysis Working Group (funded by GTEx project grants):** François Aguet[1], Shankara Anand[1], Kristin G Ardlie[1], Brunilda Balliu[8], Alvaro N Barbeira[9], Alexis Battle[10,11], Rodrigo Bonazzola[9], Andrew Brown[12,13], Christopher D Brown[14], Stephane E Castel[15,16], Don Conrad[17,18], Daniel J Cotter[19], Nancy Cox[20], Sayantan Das[21], Olivia M de Goede[19], Emmanouil T Dermitzakis[22,23,24], Barbara E Engelhardt[25,26], Eleazar Eskin[27], Tiffany Y Eulalio[28], Nicole M Ferraro[28], Elise Flynn[15,16], Laure Fresard[29], Eric R Gamazon[30,31,32,20], Diego Garrido-Martín[33], Nicole R Gay[19], Gad Getz[1,2], Aaron Graubert[1], Roderic Guigó[33,34], Kane Hadley[1], Andrew R Hamel[7,1], Robert E Handsaker[3,4,5], Yuan He[10], Paul J Hoffman[15], Farhad Hormozdiari[35,1], Lei Hou[36,1], Katherine H Huang[1], Hae Kyung Im[9], Brian Jo[25,26], Silva Kasela[15,16], Seva Kashin[3,4,5], Manolis Kellis[36,1], Sarah Kim-Hellmuth[15,16,37], Alan Kwong[21], Tuuli Lappalainen[15,16], Xiao Li[1], Xin Li[29], Yanyu Liang[9], Daniel G MacArthur[4,6], Serghei Mangul[27,38], Samuel R Meier[1], Pejman Mohammadi[15,16,39,40], Stephen B Montgomery[19,29], Manuel Muñoz-Aguirre[33,41], Daniel C Nachun[29], Jared L Nedzel[1], Duyen Y Nguyen[1], Andrew B Nobel[42], Meritxell Oliva[9,43], YoSon Park[14,44], Yongjin Park[36,1], Princy Parsana[11], Ferran Reverter[45], John M Rouhana[7,1], Chiara Sabatti[46], Ashis Saha[11], Ayellet V Segrè[1,7], Andrew D Skol[9,47], Matthew Stephens[48], Barbara E Stranger[9,49], Benjamin J Strober[10], Nicole A Teran[29], Ellen Todres[1], Ana Viñuela[50,22,23,24], Gao Wang[48], Xiaoquan Wen[21], Fred Wright[51], Valentin Wucher[33], Yuxin Zou[52]
**Analysis Working Group (not funded by GTEx project grants):** Pedro G Ferreira[53,54,55], Gen Li[56], Marta Melé[57], Esti Yeger-Lotem[58,59]
**Leidos Biomedical - Project Management:** Mary E Barcus[60], Debra Bradbury[61], Tanya Krubit[61], Jeffrey A McLean[61], Liqun Qi[61], Karna Robinson[61], Nancy V Roche[61], Anna M Smith[61], Leslie Sobin[61], David E Tabor[61], Anita Undale[61]
**Biospecimen collection source sites:** Jason Bridge[62], Lori E Brigham[63], Barbara A Foster[64], Bryan M Gillard[64], Richard Hasz[65], Marcus Hunter[66], Christopher Johns[67], Mark Johnson[68], Ellen Karasik[64], Gene Kopen[69], William F Leinweber[69], Alisa McDonald[69], Michael T Moser[64], Kevin Myer[66], Kimberley D Ramsey[64], Brian Roe[66], Saboor Shad[69], Jeffrey A Thomas[69,68], Gary Walters[68], Michael Washington[68], Joseph Wheeler[67]
**Biospecimen core resource:** Scott D Jewell[70], Daniel C Rohrer[70], Dana R Valley[70]
**Brain bank repository:** David A Davis[71], Deborah C Mash[71]
**Pathology:** Mary E Barcus[60], Philip A Branton[72], Leslie Sobin[61]
**ELSI study:** Laura K Barker[73], Heather M Gardiner[73], Maghboeba Mosavel[74], Laura A Siminoff[73]
**Genome Browser Data Integration & Visualization:** Paul Flicek[75], Maximilian Haeussler[76], Thomas Juettemann[75], W James Kent[76], Christopher M Lee[76], Conner C Powell[76], Kate R Rosenbloom[76], Magali Ruffier[75], Dan Sheppard[75], Kieron Taylor[75], Stephen J Trevanion[75], Daniel R Zerbino[75]
**eGTEx groups:** Nathan S Abell[19], Joshua Akey[77], Lin Chen[43], Kathryn Demanelis[43], Jennifer A Doherty[78], Andrew P Feinberg[79], Kasper D Hansen[80], Peter F Hickey[81], Lei Hou[36,1], Farzana Jasmine[43], Lihua Jiang[19], Rajinder Kaul[82,83], Manolis Kellis[36,1], Muhammad G Kibriya[43], Jin Billy

Li[19], Qin Li[19], Shin Lin[84], Sandra E Linder[19], Stephen B Montgomery[29,19], Meritxell Oliva[9,43], Yongjin Park[36,1], Brandon L Pierce[43], Lindsay F Rizzardi[85], Andrew D Skol[9,47], Kevin S Smith[29], Michael Snyder[19], John Stamatoyannopoulos[82,86], Barbara E Stranger[9,49], Hua Tang[19], Meng Wang[19]
**NIH program management:** Philip A Branton[72], Latarsha J Carithers[72,87], Ping Guan[72], Susan E Koester[88], A. Roger Little[89], Helen M Moore[72], Concepcion R Nierras[90], Abhi K Rao[72], Jimmie B Vaught[72], Simona Volpi[91]
**Affiliations**
1. The Broad Institute of MIT and Harvard, Cambridge, MA, USA
2. Cancer Center and Department of Pathology, Massachusetts General Hospital, Boston, MA, USA
3. Department of Genetics, Harvard Medical School, Boston, MA, USA
4. Program in Medical and Population Genetics, The Broad Institute of Massachusetts Institute of Technology and Harvard University, Cambridge, MA, USA
5. Stanley Center for Psychiatric Research, Broad Institute, Cambridge, MA, USA
6. Analytic and Translational Genetics Unit, Massachusetts General Hospital, Boston, MA, USA
7. Ocular Genomics Institute, Massachusetts Eye and Ear, Harvard Medical School, Boston, MA, USA
8. Department of Biomathematics, University of California, Los Angeles, Los Angeles, CA, USA
9. Section of Genetic Medicine, Department of Medicine, The University of Chicago, Chicago, IL, USA
10. Department of Biomedical Engineering, Johns Hopkins University, Baltimore, MD, USA
11. Department of Computer Science, Johns Hopkins University, Baltimore, MD, USA
12. Department of Genetic Medicine and Development, University of Geneva Medical School, Geneva, Switzerland
13. Population Health and Genomics, University of Dundee, Dundee, Scotland, UK
14. Department of Genetics, University of Pennsylvania, Perelman School of Medicine, Philadelphia, PA, USA
15. New York Genome Center, New York, NY, USA
16. Department of Systems Biology, Columbia University, New York, NY, USA
17. Department of Genetics, Washington University School of Medicine, St. Louis, Missouri, USA
18. Department of Pathology & Immunology, Washington University School of Medicine, St. Louis, Missouri, USA
19. Department of Genetics, Stanford University, Stanford, CA, USA
20. Division of Genetic Medicine, Department of Medicine, Vanderbilt University Medical Center, Nashville, TN, USA
21. Department of Biostatistics, University of Michigan, Ann Arbor, MI, USA
22. Department of Genetic Medicine and Development, University of Geneva Medical School, Geneva, Switzerland
23. Institute for Genetics and Genomics in Geneva (iGE3), University of Geneva, Geneva, Switzerland
24. Swiss Institute of Bioinformatics, Geneva, Switzerland
25. Department of Computer Science, Princeton University, Princeton, NJ, USA
26. Center for Statistics and Machine Learning, Princeton University, Princeton, NJ, USA
27. Department of Computer Science, University of California, Los Angeles, Los Angeles, CA, USA

28. Program in Biomedical Informatics, Stanford University School of Medicine, Stanford, CA, USA
29. Department of Pathology, Stanford University, Stanford, CA, USA
30. Data Science Institute, Vanderbilt University, Nashville, TN, USA
31. Clare Hall, University of Cambridge, Cambridge, UK
32. MRC Epidemiology Unit, University of Cambridge, Cambridge, UK
33. Centre for Genomic Regulation (CRG), The Barcelona Institute for Science and Technology, Barcelona, Catalonia, Spain
34. Universitat Pompeu Fabra (UPF), Barcelona, Catalonia, Spain
35. Department of Epidemiology, Harvard T.H. Chan School of Public Health, Boston, MA, USA
36. Computer Science and Artificial Intelligence Laboratory, Massachusetts Institute of Technology, Cambridge, MA, USA
37. Statistical Genetics, Max Planck Institute of Psychiatry, Munich, Germany
38. Department of Clinical Pharmacy, School of Pharmacy, University of Southern California, Los Angeles, CA, USA
39. Scripps Research Translational Institute, La Jolla, CA, USA
40. Department of Integrative Structural and Computational Biology, The Scripps Research Institute, La Jolla, CA, USA
41. Department of Statistics and Operations Research, Universitat Politècnica de Catalunya (UPC), Barcelona, Catalonia, Spain
42. Department of Statistics and Operations Research and Department of Biostatistics, University of North Carolina, Chapel Hill, NC, USA
43. Department of Public Health Sciences, The University of Chicago, Chicago, IL, USA
44. Department of Systems Pharmacology and Translational Therapeutics, University of Pennsylvania, Perelman School of Medicine, Philadelphia, PA, USA
45. Department of Genetics, Microbiology and Statistics, University of Barcelona, Barcelona. Spain.
46. Departments of Biomedical Data Science and Statistics, Stanford University, Stanford, CA, USA
47. Department of Pathology and Laboratory Medicine, Ann & Robert H. Lurie Children's Hospital of Chicago, Chicago, IL, USA
48. Department of Human Genetics, University of Chicago, Chicago, IL, USA
49. Center for Genetic Medicine, Department of Pharmacology, Northwestern University, Feinberg School of Medicine, Chicago, IL, USA
50. Department of Twin Research and Genetic Epidemiology, King’s College London, London, UK
51. Bioinformatics Research Center and Departments of Statistics and Biological Sciences, North Carolina State University, Raleigh, NC, USA
52. Department of Statistics, University of Chicago, Chicago, IL, USA
53. Department of Computer Sciences, Faculty of Sciences, University of Porto, Porto, Portugal
54. Instituto de Investigação e Inovação em Saúde, Universidade do Porto, Porto, Portugal
55. Institute of Molecular Pathology and Immunology, University of Porto, Porto, Portugal
56. Columbia University Mailman School of Public Health, New York, NY, USA
57. Life Sciences Department, Barcelona Supercomputing Center, Barcelona, Spain
58. Department of Clinical Biochemistry and Pharmacology, Ben-Gurion University of the Negev, Beer-Sheva, Israel
59. National Institute for Biotechnology in the Negev, Beer-Sheva, Israel
60. Leidos Biomedical, Frederick, MD, USA

61. Leidos Biomedical, Rockville, MD, USA
62. UNYTS, Buffalo, NY, USA
63. Washington Regional Transplant Community, Annandale, VA, USA
64. Therapeutics, Roswell Park Comprehensive Cancer Center, Buffalo, NY, USA
65. Gift of Life Donor Program, Philadelphia, PA, USA
66. LifeGift, Houston, TX, USA
67. Center for Organ Recovery and Education, Pittsburgh, PA, USA
68. LifeNet Health, Virginia Beach, VA. USA
69. National Disease Research Interchange, Philadelphia, PA, USA
70. Van Andel Research Institute, Grand Rapids, MI, USA
71. Department of Neurology, University of Miami Miller School of Medicine, Miami, FL, USA
72. Biorepositories and Biospecimen Research Branch, Division of Cancer Treatment and Diagnosis, National Cancer Institute, Bethesda, MD, USA
73. Temple University, Philadelphia, PA, USA
74. Virgina Commonwealth University, Richmond, VA, USA
75. European Molecular Biology Laboratory, European Bioinformatics Institute, Hinxton, United Kingdom
76. Genomics Institute, UC Santa Cruz, Santa Cruz, CA, USA
77. Carl Icahn Laboratory, Princeton University, Princeton, NJ, USA
78. Department of Population Health Sciences, The University of Utah, Salt Lake City, Utah, USA
79. Schools of Medicine, Engineering, and Public Health, Johns Hopkins University, Baltimore, MD, USA
80. Department of Biostatistics, Bloomberg School of Public Health, Johns Hopkins University, Baltimore, MD, USA
81. Department of Medical Biology, The Walter and Eliza Hall Institute of Medical Research, Parkville, Victoria, Australia
82. Altius Institute for Biomedical Sciences, Seattle, WA, USA
83. Division of Genetics, University of Washington, Seattle, WA, University of Washington, Seattle, WA, USA
84. Department of Cardiology, University of Washington, Seattle, WA, USA
85. HudsonAlpha Institute for Biotechnology, Huntsville, AL, USA
86. Genome Sciences, University of Washington, Seattle, WA, USA
87. National Institute of Dental and Craniofacial Research, Bethesda, MD, USA
88. Division of Neuroscience and Basic Behavioral Science, National Institute of Mental Health, National Institutes of Health, Bethesda, MD, USA
89. National Institute on Drug Abuse, Bethesda, MD, USA
90. Office of Strategic Coordination, Division of Program Coordination, Planning and Strategic Initiatives, Office of the Director, National Institutes of Health, Rockville, MD, USA
91. Division of Genomic Medicine, National Human Genome Research Institute, Bethesda, MD, USA

**Funding**

The consortium was funded by GTEx program grants: HHSN268201000029C (F.A., K.G.A., A.V.S., X.Li., E.T., S.G., A.G., S.A., K.H.H., D.Y.N., K.H., S.R.M., J.L.N.), 5U41HG009494 (F.A., K.G.A.), 10XS170 (Subcontract to Leidos Biomedical) (W.F.L., J.A.T., G.K., A.M., S.S., R.H., G.Wa., M.J., M.Wa., L.E.B., C.J., J.W., B.R., M.Hu., K.M., L.A.S., H.M.G., M.Mo., L.K.B.), 10XS171 (Subcontract to Leidos Biomedical) (B.A.F., M.T.M., E.K., B.M.G., K.D.R., J.B.), 10ST1035 (Subcontract to Leidos Biomedical) (S.D.J., D.C.R., D.R.V.), R01DA006227-17

(D.C.M., D.A.D.), Supplement to University of Miami grant DA006227. (D.C.M., D.A.D.), HHSN261200800001E (A.M.S., D.E.T., N.V.R., J.A.M., L.S., M.E.B., L.Q., T.K., D.B., K.R., A.U.), R01MH101814 (M.M-A., V.W., S.B.M., R.G., E.T.D., D.G-M., A.V.), U01HG007593 (S.B.M.), R01MH101822 (C.D.B.), U01HG007598 (M.O., B.E.S.).

**COI**

F.A. is an inventor on a patent application related to TensorQTL; S.E.C. is a co-founder, chief technology officer and stock owner at Variant Bio; E.R.G. is on the Editorial Board of Circulation Research, and does consulting for the City of Hope / Beckman Research Institut; E.T.D. is chairman and member of the board of Hybridstat LTD.; B.E.E. is on the scientific advisory boards of Celsius Therapeutics and Freenome; G.G. receives research funds from IBM and Pharmacyclics, and is an inventor on patent applications related to MuTect, ABSOLUTE, MutSig, POLYSOLVER and TensorQTL; S.B.M. is on the scientific advisory board of Prime Genomics Inc.; D.G.M. is a co-founder with equity in Goldfinch Bio, and has received research support from AbbVie, Astellas, Biogen, BioMarin, Eisai, Merck, Pfizer, and Sanofi-Genzyme; H.K.I. has received speaker honoraria from GSK and AbbVie.; T.L. is a scientific advisory board member of Variant Bio with equity and Goldfinch Bio. P.F. is member of the scientific advisory boards of Fabric Genomics, Inc., and Eagle Genomes, Ltd. P.G.F. is a partner of Bioinf2Bio.
